# Assessing the distinct contributions of rostral and dorsomedial prefrontal cortex to cognitive control using temporal interference brain stimulation

**DOI:** 10.1101/2024.06.06.597826

**Authors:** Johnathan S. Ryan, Boris Botzanowski, Maya Karkare, Jessica R. Kubert, Shiyin Liu, Samantha A. Betters, Adam Williamson, Negar Fani, Michael T. Treadway

## Abstract

The medial prefrontal cortex has been strongly implicated in a diverse array of cognitive functions in humans, including cognitive control and emotion regulation. Numerous studies have further proposed distinct functions for dorsomedial and rostromedial areas, but direct evidence from neuromodulation studies in healthy humans has been lacking due to the limitations of commonly used non-invasive neuromodulation techniques. Temporal interference (TI) stimulation is a recently developed technique for non-invasive deep brain stimulation that utilizes the frequency difference Δƒ between pairs of high frequency electric fields to stimulate brain regions at depth and with improved precision compared to traditional techniques. Despite its theoretical potential, however, TI applications in humans have remained limited. Here, we examined the effects of TI stimulation to dorsomedial prefrontal cortex (dmPFC) and rostromedial prefrontal cortex (rmPFC) on cognitive control. Healthy adult participants (n = 32) were recruited and administered 20 Hz Δƒ TI stimulation and 0 Hz Δƒ sham stimulation in interleaved blocks while completing two variants of the Stroop Task, a well-established paradigm intended to measure cognitive control: the Color-Word and Affective Number Stroop. During the Color-Word Stroop, we found that 20 Hz Δƒ TI stimulation of dmPFC and rmPFC relative to sham stimulation slowed down reaction times, with a significantly more pronounced slowing effect specific to incongruent trials for dmPFC stimulation as well as reduced accuracy. Importantly, effects of TI on dmPFC targets localized with fMRI differed markedly from dmPFC targeting based on a generic model, highlighting the importance of individualized targeting. For the Affective Stroop, we found that stimulation of dmPFC relative to sham stimulation facilitated increased reaction times in a valence specific-manner. This research provides novel evidence for distinct effects of neuromodulation in sub-regions of medial prefrontal cortex in healthy humans and sheds light on the strengths of TI as a non-invasive stimulation method for human cognitive neuroscience.

## Introduction

With the widespread proliferation of non-invasive neuroimaging methods, a substantial amount of human cognitive neuroscience has relied on largely correlational data for understanding cognitive processes in the human brain ^1^. The limitations of these correlational approaches have been well-documented ^2^, and the cognitive neuroscientist’s options for direct manipulations of brain circuitry in humans remains limited. One region of particular interest in this regard has been the dorsomedial prefrontal cortex (dmPFC), a brain region which is frequently engaged in a wide array of tasks and contexts as evidence by functional imaging, such that the precise nature of its function or functions remains the matter of significant debate ^3–6^. However, a common theme across studies is its role in executive control over cognitive and physical resources ^3,7^. To this end, dmPFC engagement has been observed in contexts as varied as effort-based decision-making ^8^, the effortful regulation of emotion ^9^, and cognitive control and performance monitoring ^7,4,5^, among others (see ^3^ for a comprehensive review). Given such diverse functional correlates, a unifying theory for dmPFC has been elusive, and neuromodulation approaches are valuable to better understand its functions.

To date, however, neuromodulation studies of dmPFC have been limited to either specific patient populations in which invasive brain stimulation methods can be performed as part of diagnostic assessment (e.g., epilepsy) or to more superficial aspects of dmPFC which can be targeted by noninvasive brain stimulation approaches such as transcranial magnetic stimulation (TMS) or transcranial electric stimulation (tES). Unfortunately, task-induced dmPFC activations include aspects of the dorsal anterior cingulate (dACC), mid-cingulate cortex (MCC) and pre-Supplemental Motor Area (preSMA) ^3^ that cannot be targeted by non-invasive approaches without significant stimulation of intervening tissue. It is generally accepted that TMS can only stimulate the first 6 cm of the cortex ^10^, and while tES currents can reach deeper regions, the majority of the electric field still remains in the superficial brain regions. As a result, deep brain nuclei cannot be stimulated without concurrently overstimulating the overlying regions.

Additionally, common tES techniques such as transcranial Alternating or Direct Current Stimulation (tACS/tDCS) provide limited focality due to larger electrode diameters necessary to deliver relevant brain stimulation currents without discomfort, leaving precise targeting complicated. While some prior invasive brain stimulation studies have partially corroborated neuroimaging findings–for example, demonstrating that intracranial electrode stimulation of dmPFC resulted in an “urge to act” ^11–13^;– the development of more effective non-invasive approaches to stimulation of dmPFC and other cingulate regions is needed.

In addition to a central role in cognitive control, the dmPFC has increasingly been implicated in processes related to emotion and emotion regulation ^9,14^. Indeed, while initial theoretical work informed by the first decade of neuroimaging studies postulated a dorsal/ventral divide of the medial wall such that dmPFC and surrounding areas were restricted to “cold” cognitive processes ^15^, more recent work has found robust evidence for dmPFC processing of affective stimuli, particularly those related to negative affect ^16,17^ and pain ^18^, as well as pleasure^19^. However, the effects of causal manipulations to this region on cognitive-emotion interactions remain unclear.

In attempting to overcome the limitations of other tES methods, a new non-invasive brain stimulation technique, temporal interference (TI), has emerged ^20–22^. TI utilizes temporally interfering electric fields to stimulate deep brain regions with great precision, offering the opportunity to garner novel insight of targeted structures and related mechanisms. The temporal interference technique is based on the use of two or more differing high-frequency signals (> 1000 Hz) which are conventionally deemed to not individually affect neurons ^23,24^. When these waves overlap at a spatial point, their electric fields (E-Field) combine through the principle of superposition, creating what is known as an amplitude-modulated (AM, sinusoidal envelope) or pulse-width-modulated (PWM, square wave modulation) ^25^ signal at a frequency equal to the difference (Δƒ) between the two source signals (|ƒ1 – ƒ2| = Δƒ). The high frequencies used in TI allow for a deeper penetration of electric fields, and, most importantly, this technique seems to offer the ability to stimulate specific deeper regions without activating surrounding/overlaying brain tissue.

In the current study, we sought to leverage TI to better understand the effects of cognitive control in the context of affective and non-affective stimuli using two versions of the classic Stroop task: the Color-Word Stroop ^26^ (**Fig. 1A**) and the Affective number Stroop ^27^ (**Fig. 1B**). Using MRI-guided functional localization, we evaluated the differential effects of excitatory TI stimulation (Δƒ=20 Hz) vs. sham stimulation (Δƒ=0) on Color-Word Stroop and Affective Stroop task performance when targeting the dmPFC, specifically examining the Stroop effect, which contrasts trials that assay effects of interference (trials in which the stimulus word color or number of digits displayed is incongruent with the actual color or number displayed for the Color-Word Stroop and Affective Stroop, respectively) vs facilitation (trials in which the stimulus color or stimulus number of digits displayed is congruent with the actual color or number displayed, for the Color-Word Stroop and Affective Stroop, respectively). The dmPFC target region was defined individually using single-subject fMRI contrast maps of incongruent > congruent trials during the Color-Word Stroop. For a comparison site, we also targeted an individualized region of rostromedial prefrontal cortex (rmPFC) that was more active during congruent trials relative to incongruent trials during the Color-Word Stroop.

**Figure 1:**
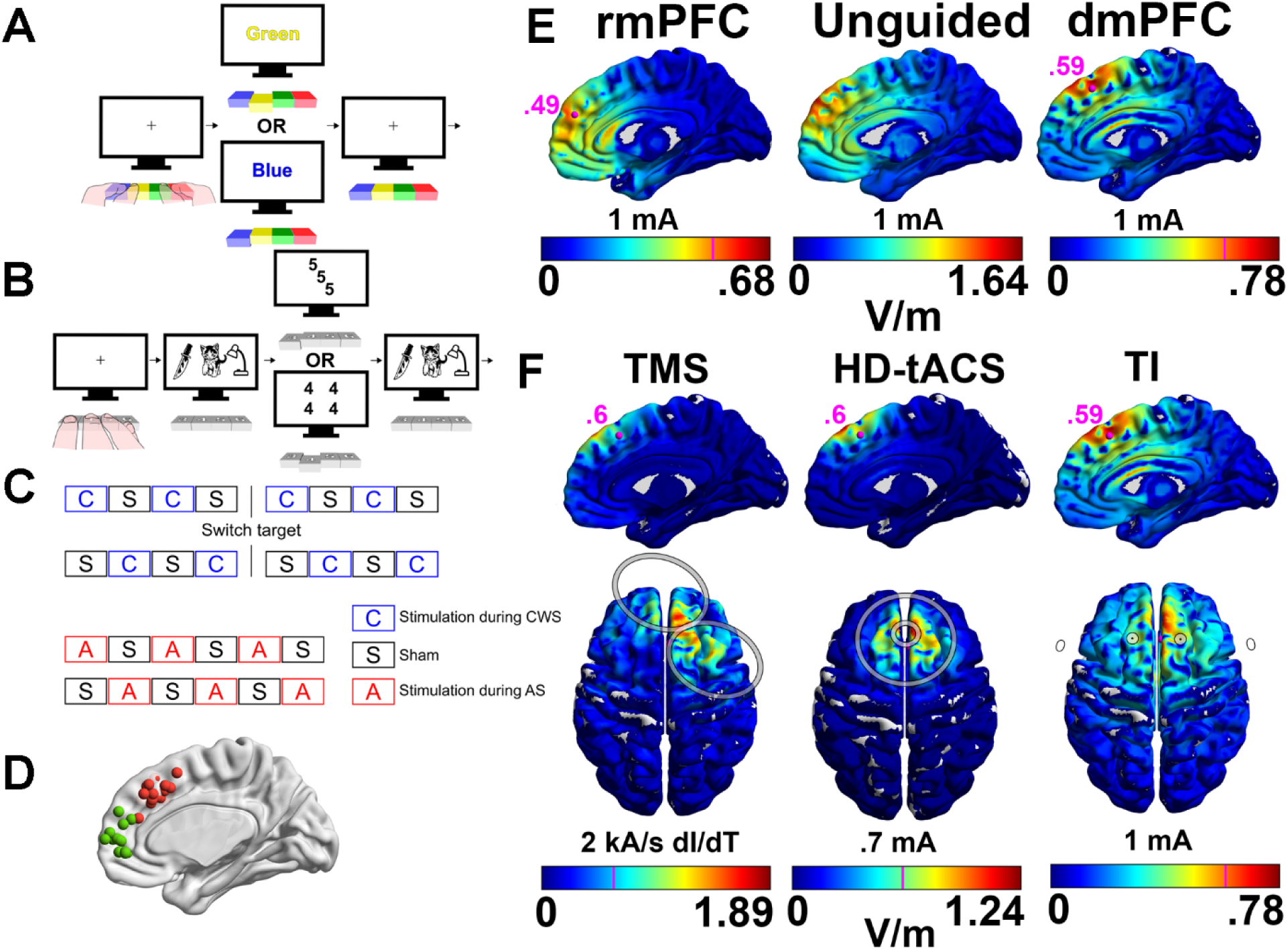
Task design, experimental procedures and fMRI results. **A-B.** During separate fMRI scanning and TI stimulation visits using 20 Hz, participants performed Stroop tasks. For the Color-Word Stroop, they were instructed to use their left and right index fingers to press the adequate key on a keyboard to select their answer. For the Affective Stroop they were instructed to use their right index, middle, ring and little finger to press the adequate key on a keyboard to select their answer. **C**. Block-design for TI vs sham stimulation. Participants underwent 6 blocks (for unguided dmPFC targeting) or 8 blocks (for MRI-guided dmPFC/rmPFC targeting) of Color-Word Stroop tasks, and 6 blocks of Affective Stroop tasks. After the first 4 blocks of Stroop tasks, the stimulated target was changed from dmPFC to rmPFC and vice versa for the sessions with individualized targeting. The order of stimulation, sham vs stimulation, and Affective Stroop (AS) vs Color Word Stroop (CWS) was randomized. **D.** Depiction of individually localized (MRI-guided) stimulation targets for dmPFC (red) and rmPFC (green). **E.** Example electric fields resulting from 20 Hz Δƒ TI using representative electrode placements for unguided rmPFC, unguided dmPFC and MRI-guided dmPFC targets. Example Individual target shown with pink sphere and volts/meter at target shown in pink. **F.** Example of simulated electric fields and resulting volts/meter stimulation resulting from different stimulation methods and coil/electrode placements (gray) contrasting focality of TI vs TMS and high-definition tACS.

In addition to seeking to determine the effects of TI on dmPFC function during cognitive control within emotional and non-emotional contexts, our study was also designed to address several open questions about the nature of TI stimulation. First, we sought to compare the effects of functionally-localized targeting (using fMRI) vs. unguided targeting (using a normative brain model to guide electrode placements). Second, the extent to which carry-over effects, that is the continued impact of stimulation immediately after stimulation has been stopped, is unknown. As such, we employed a blocked design with active stimulation (TI On (Δƒ>0)) vs sham stimulation (TI Off (Δƒ=0) and analyzed our results by comparing TI On vs TI Off blocks as well as effect of cumulative TI stimulation time.

## RESULTS

### Neuroimaging Results

As expected, significantly greater activation in the dmPFC was observed during incongruent compared to congruent Color-Word Stroop trials (*p* = 8.6e-4; See Supplemental Materials). Importantly, the area of dmPFC showing the strongest differences varied significantly across subjects (**Fig. 1E**), highlighting the importance of individual targeting. Additional results from neuroimaging are included in the supplement.

### TI Block Analyses: Color-Word Stroop

For all behavioral analyses, we used linear-mixed effects models to investigate the effects of TI stimulation on task reaction time (RT) and accuracy. Initial analyses explored the effect of block type (TI On or Off). For Color-Word Stroop, a linear mixed effect model revealed a main effect of condition (*t* = 9.774, *p* < 6.25e-21, *p_FDR_* = 6.29e-21), and stimulation site (dmPFC vs. rmPFC: *t* = 2.508, *p* = .0122, *p_FDR_* = .0376). There was a significant block type × site interaction (unguided dmPFC vs. rmPFC: *t* = –2.073, *p* = .0382, *p_FDR_* = .0929) but the main effect of block type was non-significant (*t* = 0.493, *p* = .6219, *p_FDR_* = .755), with unguided dmPFC showing a reduction of the Stroop effect and rmPFC showing a potentiation of the Stroop effect. Full model results can be found in Supplemental Materials.

One explanation for this null block effect may be carry-over effects from stimulation that impacted the target function even during sham stimulation. To examine this possibility, we conducted two sets of analyses. The first was a between-subjects comparison of individuals who performed the Color-Word Stroop task first (*n* = 11). This analysis showed a nominally significant effect of block type (TI On vs Off) (*t* = 2.61, *p* = .0439, *p_FDR_* = .298) and trend-level block × site interactions (dmPFC vs. rmPFC: *t* = –2.06, *p* = .09, *p_FDR_* = .298; unguided dmPFC vs rmPFC: *t* = –2.183, *p* = .0759, *p_FDR_* = .298), though these effects did not survive FDR correction. Full model results can be found in Supplemental Materials.

As a second test, we also did a within-subject comparison for the same set of 11 subjects who completed the Color-Word Stroop task first and examined their performance during the first TI On Block vs the first TI Off Block. The rationale for this analysis was that carry-over effects would be relatively minimal after just a single block of TI On, providing greater power to detect effects. Consistent with this hypothesis, we observed a nominal effect of block type (TI On or TI Off) (*t* = 1.997, *p* = .046; *p_FDR_* = .148) and a block type × site interaction (dmPFC vs. rmPFC: *t* = –3.41, *p* = .0007; *p_FDR_ =* .004; unguided dmPFC vs. rmPFC: *t* = –4.553, *p* = 5.99e-6, *p_FDR_* = 9.58e-5). Moreover, these effects were strongest in individuals who began with TI Off in their first block and then completed the task with TI On, and the difference between TI On and Off was attenuated in subsequent task runs, both consistent with carry-over effects (see Supplemental Materials). While these results were only nominally significant due to being performed in a subset of participants, they nevertheless support the hypothesis that TI induces carryover effects. Full model results, and an additional model including stimulation order, can be found in Supplemental Materials.

### TI Block Analyses: Affective Stroop

A similar pattern of results emerged for Affective Stroop. A linear mixed effect model revealed a main effect of condition (*t* =9.475, *p* = 3.83e-21, *p_FDR_* = 6.51e-20), and a marginal stimulation site × block type interaction (dmPFC vs. unguided dmPFC: *t* = –1.841, *p* = .066, *p_FDR_* = .279) with the dmPFC site showing an increase in overall response time during the TI block and the unguided dmPFC site showing a decrease in overall response time. The distractor stimuli (Aversive, pleasant, neutral) did not show significant valence effects (neutral vs. pleasant: *t* =1.575, *p* = .1153, *p_FDR_* = .368; neutral vs. aversive: *t* =1.08, *p* = .2803, *p_FDR_* = .622; aversive vs. pleasant: *t* =.475, *p* = .6347, *p_FDR_* = .922). Full model results can be found in Supplemental Materials.

As with the Color-Word Stroop, we considered the influence of carry-over effects by examining participant’s performance during the first TI On Block vs the first TI Off Block for individuals who ran the Affective Stroop task first (*n* = 9), which showed a three-way block × site × distractor interactions (neutral vs. aversive: *t* = 1.974, *p* = .0494, *p_FDR_* = .317; pleasant vs. aversive: *t* = 2.014, *p* = .0450, *p_FDR_* = .317), and a four-way block × condition × site × distractor interaction (pleasant vs. aversive: *t* = –2.056, *p* = .0408, *p_FDR_* = .317). To capture additional within subject effects, analysis of the first pair of TI On/Off blocks revealed an effect of site (dmPFC vs. unguided dmPFC: *t* = 2.221, *p* = .045; *p_FDR_* = .208), a block type × site interaction (dmPFC vs. unguided dmPFC: *t* = –2.67, *p* = .0008, *p_FDR_* = .085) and a three-way block × distractor × site interaction (neutral vs. aversive: *t* = 2.49, *p* = .0131, *p_FDR_* = .105). Full model results can be found in Supplemental Materials.

### TI Cumulative Stimulation Time: Color Word Stroop Reaction Time

Given the evidence for stimulation carry-over effects from TI On blocks to TI Off blocks, we next considered a predictor variable representing cumulative TI stimulation time for each target. Control analyses were performed to rule out the confound of fatigue or practice effects that might also increase monotonically along with cumulative TI stimulation (see Control Analyses below). Additionally, because cumulative stimulation time would differ as a function of task run order (e.g., whether one performed the Color-Word Stroop task at the beginning or end of the session), we included run-order as a nuisance co-variate for all analyses. For each task, we first conducted an omnibus model including task condition, cumulative TI stimulation time, and target stimulation site. For Color-Word Stroop (**Fig. 3A**), this model revealed main effects of condition (congruent vs. incongruent: *t* = 5.851, *p* = 5.04e-09, *p_FDR_* = 3.43e-8), stimulation time (*t* = 3.070, *p* = .002, *p_FDR_* = .009), and stimulation site (dmPFC vs. rmPFC: *t* = 2.607, *p* = .009, *p_FDR_* = .031; dmPFC vs. unguided dmPFC: *t* = 2.869, *p* = .007, *p_FDR_* = .026; unguided dmPFC vs. rmPFC: *t* = –2.365, *p* = .0234, *p_FDR_* = .061). Stimulation time × site interactions revealed significant RT reductions over stimulation time during unguided dmPFC targeting relative to the rmPFC (*t* = 5.489, *p* = 4.31e-8, *p_FDR_* = 2.44e-7) and dmPFC (*t* = 6.117, *p* = 1.06e-9, *p_FDR_* = 9.01e-9) targets, which saw increased RTs as stimulation time increased; dmPFC vs. rmPFC did not differ significantly (*t* = –1.612; *p* = .107, *p_FDR_* = .214). Individual models split by site show effects of stimulation time on the rmPFC site (*t* = 2.612, *p* = .009) and unguided dmPFC site (*t* = –5.747, *p* = 1.01e-08) and a marginal effect on the dmPFC site (*t* = 1.707, *p* = .088). Three-way condition × stimulation time × stimulation site interactions revealed unguided dmPFC targeting led to reduced RTs as stimulation time increased, and differed significantly from dmPFC (*t* = –3.166, *p* = .0016) and marginally to rmPFC targeting (unguided dmPFC vs. rmPFC; *t* = 1.77, *p* = .076), which both showed increased RTs with increased stimulation time (dmPFC vs. rmPFC did not meet significance, *t* = –1.475; *p* = .14). This difference in RT when comparing dmPFC and unguided dmPFC was similar for both conditions (congruent: *t* = –4.438, *p* = 1.02e-5; incongruent: *t* = –5.716, *p* = 1.35e-8) but when comparing dmPFC to rmPFC, the difference in RT was stronger for incongruent trials (congruent: *t* = –1.938; *p* = .053; incongruent: *t* = –3.353, *p* = 8.1e-4) trials. Individual models split by target site revealed stimulation time effects for rmPFC (*t* = 2.639, *p* = .008), unguided dmPFC (*t* = –5.752, *p* = 9.78e-9), and marginally for dmPFC (*t* = 1.711, *p* = .087).

**Figure 2:**
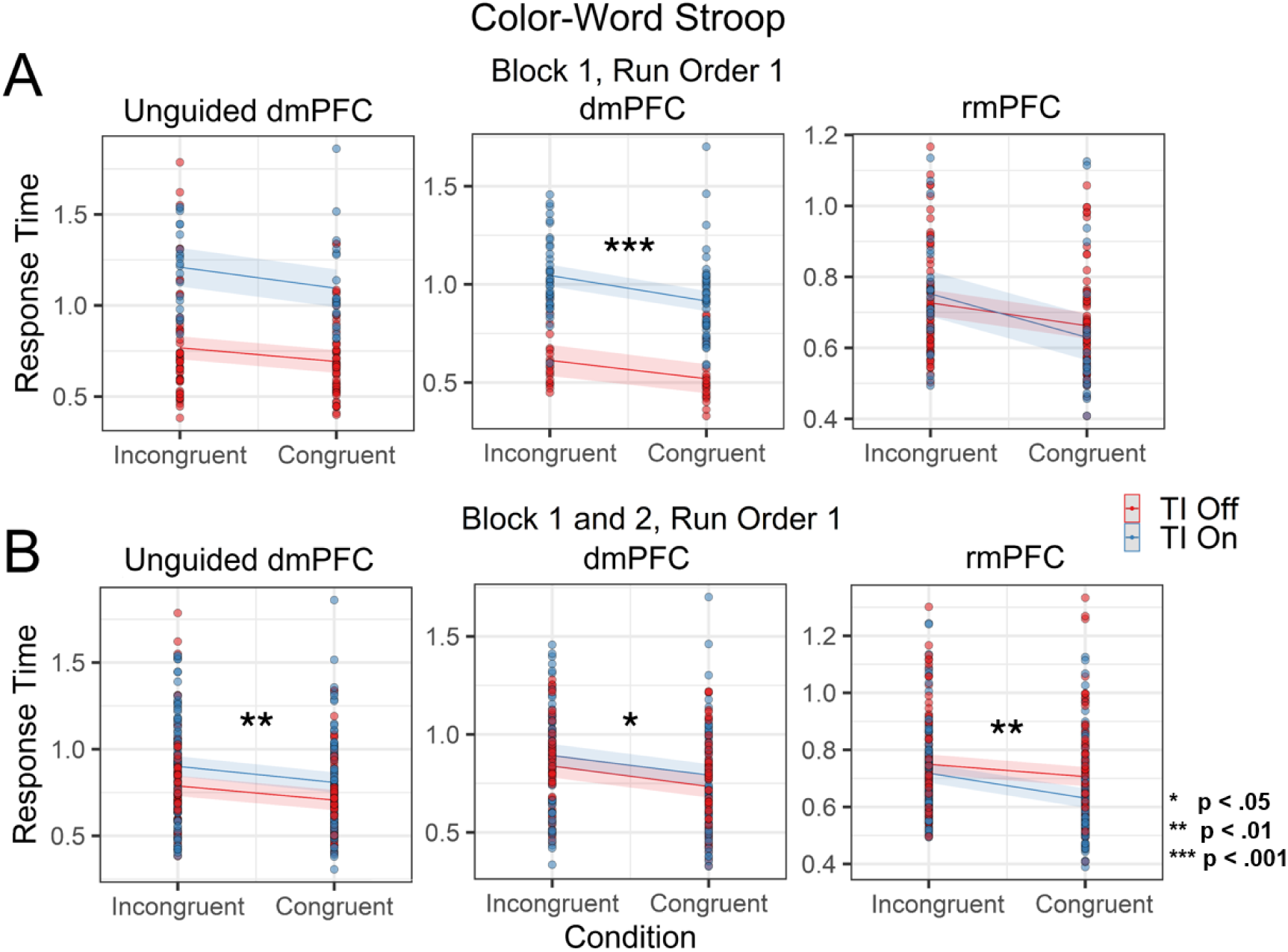
Block 1 and Block 2, Run1 Comparisons. TI effects during **A**. Block 1 and **B**. Block 1 and 2 when Color-Word Stroop was administered first, thereby limiting the possible contribution of carry-over effects. Data points displayed are individual trials.

**Figure 3:**
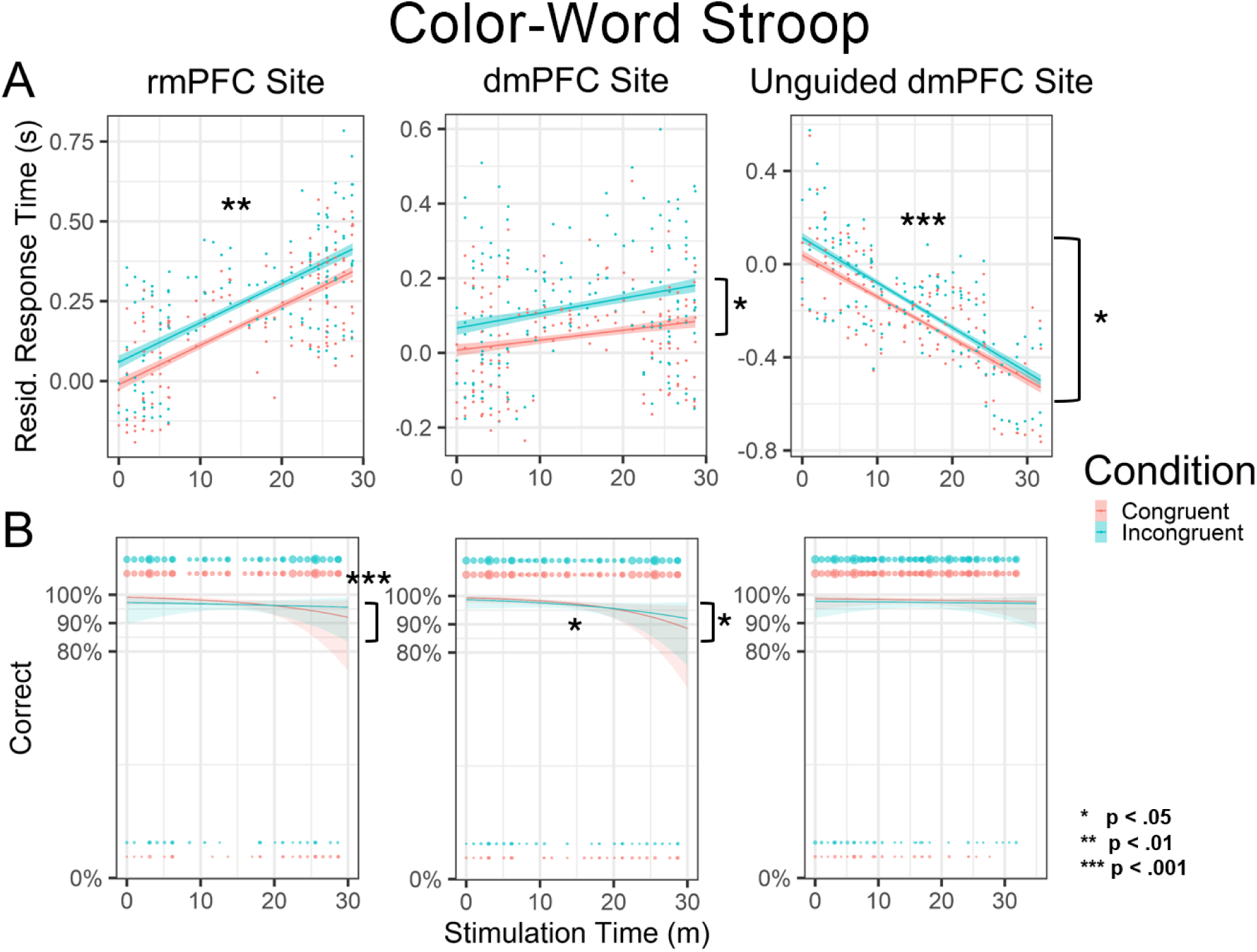
Effects of cumulative stimulation time on reaction time and accuracy during the Color Word Stroop. **A.** Increasing TI stimulation time was associated with site-specific effects on reaction times and the Stroop effect. **B.** Increasing TI stimulation time was associated with site-specific effects on accuracy. Circle size indicates frequency of correct (top) or incorrect (bottom) trials at each unique time point.

To further unpack how TI stimulation time effects differed by target site, we next tested condition × stimulation time interactions within these separate models for each of the three targets. These models found that increasing TI stimulation time led to an increased Stroop effect for the dmPFC (*t* = 2.388, *p* = .017) and a decreased Stoop effect for unguided dmPFC (*t* = – 2.267, *p* = .023); importantly, the Stroop effect did not seem to be affected by stimulation to rmPFC (interaction was non-significant (*t* = .160, *p* = .873). Individual models split by condition and site revealed effects for rmPFC congruent (marginal; *t* = 1.847, *p* = .065), rmPFC incongruent (*t* = 2.016, *p* = .044), unguided dmPFC congruent (*t* = –3.783, p = 1.62e-4), unguided dmPFC incongruent (*t* = –5.630, *p* = 2.18e-8), and congruent dmPFC (marginal; *t* = 1.909, *p* = .056) trials; stimulation time effects were not significant for incongruent dmPFC trials (*t* = 1.486, *p* = .137).

### TI Cumulative Stimulation Time: Color-Word Stroop Accuracy

A similar model exploring stimulation time effects on accuracy (**Fig. 3b**) revealed a main effect of condition (*z* = –2.301; *p* = .021, *p_FDR_* = .123), stimulation time (*z* = –2.890, *p* = .004, *p_FDR_* = .035), run order (run order 1 vs. 2: *z* = 1.884, *p* = .06, *p_FDR_* = .142; run order 1 vs. 3: *z* = 1.983, *p* = .047, *p_FDR_* = .142) and a stimulation × condition interaction (*z* = 2.112, *p* = .035, *p_FDR_* = .125). Consistent with effects on RT, individual models split by site showed significant condition by stimulation time effects for dmPFC (*z* = 2.117, *p* =.034) and rmPFC sites (*z* = 3.306; *p* = .0009); unguided dmPFC site effects did not reach significance (*z* = .324, *p* = .746). Interestingly, for the dmPFC, this effect was driven by a reduced accuracy that was significant only for incongruent trials (*z* = –1.961, *p* = .05), suggesting that MRI-Guided dmPFC TI stimulation impaired both speed and accuracy of incongruent trials relative to congruent trials. For rmPFC, the effect of cumulative stimulation was specific to congruent trials (*z* = –2.451, *p* = .014).

### TI Cumulative Stimulation Time: Affective Stroop Reaction Time

A similar model was conducted with the Affective Stroop, with the addition of distractor type (aversive, pleasant, neutral). Unlike the Color-word Stroop, the Affective Stroop did not include the rmPFC control site, so only two stimulation sites, guided and unguided dmPFC were used. As both sites targeted the same region and there was no omnibus interaction with site, analyses were collapsed across both dmPFC sites. This model revealed a main effect of condition (*t* = 8.197, *p* = 3.00e-16, *p_FDR_* = 3.06e-15), stimulation time (*t* = –11.044, *p =* 4.53e-28, *p_FDR_* = 1.16e-26), run order (run order 1 vs. 2: *t* = 3.401, *p* = .002, *p_FDR_* = .015; run order 2 vs. 3, *t* = 2.716; *p* = .002), and a marginal main effect of distractor valence (pleasant vs. aversive: *t* = –1.803; *p* = .071, *p_FDR_* = .28) (**Fig. 4a**). Stimulation time × distractor valence interactions showed greater RT slowing as stimulation time increased during neutral distractor trials relative to pleasant distractor trials (*t* = 2.812, *p* = .005, *p_FDR_* = .028) and aversive trials relative to pleasant trials (*t* = 1.953, *p* = .051, *p_FDR_* = .259); neutral vs. aversive did not reach significance (*t* = .817, *p* = .414, *p_FDR_* = .813). Individual models split by valence showed significant effects of stimulation time on pleasant (*t* = –6.403, *p* = 1.91e-10), aversive (*t* = –6.489, *p* = 1.11e-10) and neutral distractor trials (*t* = –5.841, *p* = 6.18e-09).

**Figure 4:**
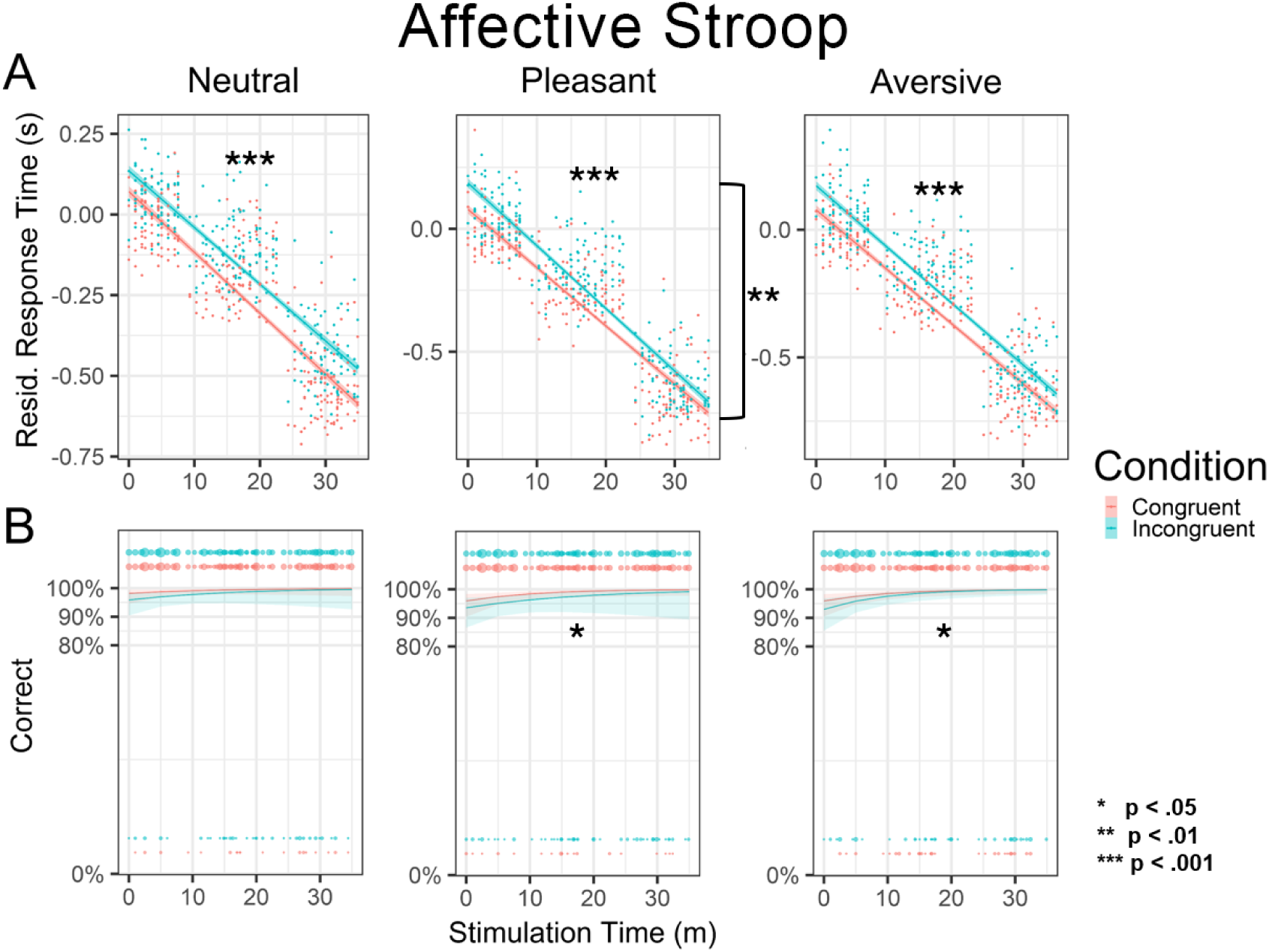
Effects of cumulative stimulation time on reaction time and accuracy during the Affective Stroop. **A.** Increasing TI stimulation time was associated with a general speeding of RT across all distractor-valence conditions but led to a reduced Stroop effect for pleasant and Aversive distractors relative to neutral. **B.** Increasing TI stimulation time was associated with improved accuracy across distractor-valence conditions.

Three-way condition × stimulation time × valence interactions revealed greater RT slowing over stimulation time during neutral trials relative to pleasant (*t* = –2.929, *p* = .003) and aversive trials (*t* = –1.667, *p* = .096); pleasant trials did not differ significantly from aversive trials (*t* = –1.253, *p* = .210). Similarly, an additional model comparing emotional vs. neutral distractors shows a similar condition × stimulation time × distractor type interaction (t = 2.677, p = .007). Individual models split by valence show a significant stimulation time × condition effect during pleasant trials (*t*=-2.7, *p* = .007), with incongruent trials showing significantly reduced RT, suggesting a reduction of the Stroop Effect. These effects were not significant for aversive (*t* = –.882, *p* = .378) or neutral (*t* = 1.452, *p* = .147) distractor trials. Additional models split by condition and distractor valence revealed stimulation time effects for neutral congruent (*t* = – 2.88, *p* = .004), neutral incongruent (*t* = –4.52, *p* = 6.88e-6), pleasant congruent (*t* = –4.306, *p* = 1.84e-5), pleasant incongruent (*t* = –5.172, *p* = 2.81e-7), aversive congruent (*t* = –2.88, *p* = .004) and aversive incongruent trials (*t* = –5.023, *p* = 6.07e-7).

### TI Cumulative Stimulation Time: Affective Stroop Accuracy

A similar model exploring stimulation time effects on accuracy (**Fig. 4b**) revealed a marginal effect of condition (*z* = –1.682; *p* = .093, *p_FDR_* = .293), stimulation time (*z* = 2.68, *p* = .007, *p_FDR_* = .035) and run order (run order 1 vs. 2: *z* = –3.095, *p* = .002, *p_FDR_* = .012; run order 1 vs. 3: *z* = – 3.787, *p* = .0002, *p_FDR_* = .001; run order 2 vs. 3: *z* = –2.815, *p* = .005). Additionally, there was an interaction of distractor valence × stimulation time (aversive vs. pleasant: *z* = 1.955, *p* = .0509, *p_FDR_* = .193), suggesting increased stimulation time led to greater accuracy for pleasant relative to aversive distractor trials. Individual valence models showed improved accuracy with increased stimulation time on pleasant (*z* = 2.119, *p* = .034) and aversive distractor trials (*z*= 2.483, *p* = .013); neutral trials were non-significant (*z* = 1.467, p =.142). Condition × stimulation time was not significant for neutral (*z* = –.32, *p* = .749), pleasant (*z* = –1.59, *p* = .112), and aversive distractor trials (*z* = .410, *p* = .681). Models split by valence and condition showed improved accuracy with increased stimulation time for congruent aversive distractor trials (*z* = 3.292, *p* =.001) and marginal effect for congruent neutral distractor trials (*z* = 1.648, *p* = .099) but was non-significant for congruent pleasant (*z* = 1.424, *p* = .155), incongruent pleasant (*z* = 1.127, *p* = 0.26), incongruent neutral (*z* = .679, *p* = .497), and incongruent aversive trials (*z* = .948, *p* = .343).

### Control Analyses

#### Inclusion of On/Off Block Variable

Inclusion of a block variable into the stimulation time models did not result in any significant main effects of block. Taken together, results from these models suggested that TI effects were heavily dependent on the total amount of stimulation time received by each region, consistent with prior work from other tES modalities suggesting carry-over entrainment effects that may persist for minutes to hours ^28^. Results from this model emphasize greater effects from overall stimulation time compared to block type, that stimulation altered Stroop processing over time, and with additional site effects in Color-Word Stroop.

#### Controlling for Passage of Time, Fatigue and Practice Effects

Because stimulation time was potentially confounded by the mere passage of time, we performed several additional analyses to rule out practice and fatigue effects. First, we tested the effect of trial number on RT performance for both tasks. For Color-Word Stroop, we found that trial number did not predict RT (*t* = .691, *p* =.489), accuracy (*t* =.012, *p* = .99), or the Stroop Effect (*t* = –.749, *p* =.454). Additionally, when trial number was included in our models with stimulation time, we continued to observe interactions with target site and condition (see Supplement). For the Affective Stroop, we did see a main effect of trial number on reaction time (*t* = –7.438, *p* = 1.17e-13) and accuracy (*z* = 3.332, *p* = .0009), but continued to observe similar interactions with distractor valence for response time. With the inclusion of trial number, stimulation effects on accuracy for the omnibus were non-significant. Full models included in supplemental methods.

## Discussion

In the current study, we sought to reveal effects of TI on human cognitive control by modulating the function of multiple sites of prefrontal cortex. TI is a relatively new non-invasive stimulation method, and its application to human cognitive neuroscience remains in nascent stages. Here, we demonstrated site-specific effects of TI stimulation performance during task conditions requiring varying levels of cognitive control and emotion regulation. In particular, the effects of dmPFC stimulation were consistent with prior results using other forms of non-invasive stimulation, but the use of TI also allowed us to easily conduct within-subject comparisons to a control site (rmPFC) with a level of focality likely not possible with other non-invasive methods (see **Fig1 F**). These results highlight the focality of TI and its promise as a tool for cognitive neuroscience.

During the Color-Word Stroop task, we observed differential effects of stimulation time on task performance as a function of site. When using MRI-guided targeting of the dmPFC, we found that the “Stroop Effect” (i.e, the difference in RT between congruent and incongruent trials) was increased during stimulation, and that this effect was specific to increased speed for incongruent trials. Similar effects have been found when using TMS with a double-cone coil that can penetrate somewhat deeper into cortex as compared to a conventional butterfly coil^29,30^, though unlike TI, will still stimulate more superficial tissue as well (see **Fig. 1F**).

Another critical advantage of TI was the ability to compare the effects of dmPFC stimulation to a control rmPFC region. In contrast to dmPFC, stimulation to rmPFC resulted in a general increase in RT without any condition-specific effects, resulting in no change to the Stroop-effect. These data are consistent with the proposed role for dmPFC as being particularly involved in cognitive control. Given that our protocol used a putatively excitatory frequency within the mid-beta range ^31^, our targeting of dmPFC could be viewed as enhancing engagement of this region during incongruent trials that have already been found to increase dmPFC activity in prior studies as well as the current sample (See **Fig. 1**). While precise functions of dmPFC are debated, there is general consensus that activity is increased under conditions of surprise, error, decision conflict and decision-difficulty ^3–6^, all of which have in turn been robustly associated with longer response times. This pattern of results is therefore consistent with the observation that TI stimulation selectively enhanced dmPFC engagement during incongruent trials.

It is noteworthy that this pattern of results was not observed when dmPFC was targeted without using MRI guidance. As TI becomes a more widely-used technique, a critical question will be whether individual localized targets will be necessary. Indeed, many TI studies published in humans to date have not used MRI-guided targeting^32–34^. In the present study, we observed that our unguided targeting of dmPFC resulted in a general speeding of reaction time and a reduction of Stroop the effect. While we do not have MRI data on these participants, based on simulations (see Supplemental Materials), we believe that our unguided electrode placements likely reached a broader swatch of supplemental motor area (SMA) and pre-SMA, which may have elicited stronger motor responses. It is also important to mention that the electric field orientation for our unguided placements was transverse as opposed to sagittal for our MR-guided placements, which may led to the stimulation of more supplementary and primary motor areas, thereby speeding motor responses ^32,35^. While we cannot be certain as to why the unguided dmPFC results were so different, they do highlight the importance of MRI-guided targeting for future TI studies.

During the Affective Stroop, cumulative TI stimulation was associated with speeding reaction time across all three distractor valence conditions. Of note, while we did not observe a significant difference between the MRI-guided and unguided dmPFC sites, these effects were generally stronger for the unguided dmPFC, which similarly showed speeded reaction times during the Color-Word Stroop. Interestingly, we also observed specific effects as a function of distractor valence, with greatest reductions of reaction time for incongruent trials during emotion (pleasant or negative) stimuli as compared to neutral stimuli. Similar results were found for accuracy, highlighting the role for affective stimuli to influence cognitive control in dmPFC. Overall, effects of TI on stimulation to guided or unguided dmPFC during the affective stroop were more similar to the effects of unguided dmPFC stimulation during the Color-word stroop. Given that this effect was present across both tasks for the unguided placement that reached more preSMA and SMA areas, it suggests that stimulation hear may help motor areas override the influence of affective distractors, consistent with prior stimulation studies suggesting that stimulation to this area may promote an “urge to act” ^11–13^.

Our results also shed some light on the possible mechanisms underlying TI stimulation. While a number of studies have already described the effects of TI stimulation on rodents, particularly on the central and peripheral nervous systems ^20,25,36–41^, relatively few human studies have been conducted ^32–34,42,43^. Rodent studies have suggested robust changes in firing rates as a function of TI “on” ( Δƒ >0) vs TI “off” (Δƒ = 0) ^20^. However, the biophysics of stimulating the much smaller rodent brain differ substantially from those in humans. Moreover, these rodent studies have used voltages that are an order of magnitude larger than what may be safely applied in humans ^44–46^. An alternative proposal has that TI may exert influence on cognitive processes through oscillatory entrainment effects, similar to other tES approaches ^47^. Neuronal oscillations within different frequencies bands play a critical in coordination of local and distal information processing ^48,49^. Supporting this proposed mechanism, one recent study in non-human primates found that TI exhibited clear effects on neuronal oscillations without changes in firing rate, as measured by implanted electrodes ^44^. These data suggest that future TI studies will need to carefully assess and control for carry-over effects when using single-session designs.

### Limitations

While the present study has a number of strengths, it also suffers from several limitations. First and foremost, our intended block design did not reveal TI effects. The failure of our block design is likely due to pronounced carry-over effects, which have been observed with other tES methods. Indeed, carryover effects have also been measured using EEG post-tACS that may persist up to an hour or longer from a single stimulation ^50,51^.

While we did observe significant effects using a measure of cumulative TI stimulation, this variable was potentially confounded by the mere passage of time. We performed a number of control analyses to rule out this possibility, and show that effects of stimulation time largely remained even when controlling for trial number. Moreover, for the Color-Word Stroop, trial number was not a significant predictor of speed or accuracy, suggesting that the observed effects with cumulative stimulation time were indeed due to TI stimulation. Nevertheless, it is possible that the passage of time exerts non-linear effects that our trial number regressor does not fully capture. Future studies should seek to estimate the duration of TI carry-over effects and account for their duration in the performance of TI On vs Off blocks of trials.

A second limitation is that we were not able to examine effects of TI stimulation to rmPFC during the Affective Stroop. While we successfully administered TI without any serious adverse events, emphasizing its safety and tolerability (see Supplement), we nevertheless were not able to target all regions due to duration of the TI sessions.

## Materials and Methods

### Participants and Study Design

Thirty-five healthy volunteers (17 (57%) female; Mean age=24.18 ± 5.9 years) were recruited from the general population using flyers posted near Emory University and through social media advertisements. Participants completed two study visits. In the first visit, participants arrived at the TReADLab in the psychology building at Emory University. Participants provided informed consent and were then escorted to the Facility for Education and Research in Neuroscience (FERN) to undergo structural MRI, and complete fMRI versions of the Color Word Stroop and the Affective Stroop task utilizing an MRI compliant button box with stimuli being presented using a back projection mirror. In the second session, participants completed the same tasks during a single-blind stimulation session with active and Sham TI. Results from a third task collected during the TI visit will be published separately. Prior to participation, all volunteers underwent screening for potential contraindications for TI or fMRI. One participant did not complete the fMRI while two participants did not complete TI (see Supplemental Methods). For Color-Word Stroop, one participant was removed for not exhibiting the anticipated Color-Word Stroop effect and one participant was removed (for non-compliance during the Color-Word Stroop task only). For Affective Stroop, two participants targeted at the rmPFC were not included. For both tasks, a final sample of 30 participants were utilized for analysis, with two unique participants between tasks. All procedures were approved by the Emory University IRB and all participants provided voluntary informed consent prior to their involvement in the study. Participants received a fixed compensation rate of $30 per hour.

### Task design

Participants engaged in two tasks: the Color-Word Stroop task and the Affective Stroop task. In the Color-Word Stroop task, participants observed a series of words displayed in different colors, with each word appearing for 1.5 seconds (**Fig. 1A**). Their goal was to determine as fast as possible the color of the word using specific keyboard keys. Stimuli could be congruent, meaning the word’s color matched the word itself (e.g., “BLUE” written in blue), or incongruent, indicating that the word’s color did not match the word (e.g., “BLUE” written in yellow). Each block was composed of 44 trials. In the Affective Stroop task, participants were shown a series of varying numerical digits. The appearance of these digits was preceded and followed by the brief appearance of a distractor with either a negative, pleasant, or neutral emotional valence (**Fig. 1B**). Participants were required to press a designated key corresponding to the number of digits displayed. Stimuli could be congruent (where the number of numbers matched the value of the numbers) or incongruent (where the number of numbers did not match the value of the numbers). Each block was composed of 33 trials.

### TI protocol

We applied TI using two DS5 Isolated Bipolar Constant Current Stimulators (Digitimer, Hertfordshire, UK). A Keysight waveform generator (Keysight Technologies Inc., Santa Rosa, CA) was employed to create a stimulation signal delivered to each stimulator. To generate a TI envelope of 20 Hz, we set frequencies to 5000 Hz (f1) for one stimulator and 5020 Hz (f2) for the second stimulator, resulting in a TI envelope frequency of 20 Hz (f1-f2 = Δf). The amplitude for each pair was standardized at 1 mA resulting in an estimated e-field 2 mm sphere average of .600 V/m for the dmPFC target and .635 V/m for the rmPFC target (see: Electromagnetic field computation for simulation methodology). Sham stimulation consisted in applying two envelope-free carriers, i.e. 5000 and 5000 Hz, at the same coordinates and amplitude as the TI. We used electrode from an EGI Geodesic Sensor Net (Electrical Geodesics Inc., Eugene, OR) to deliver TI. A conductive gel paste was applied to the stimulation electrodes to facilitate current transfer and improve participant comfort. For MRI-guided stimulations, electrode placement was determined based on Sim4Life^52^ TI simulations and a Brainsight TMS-MRI co-registration system (Rogue Research, Quebec, Canada) with a Polaris stereo camera. Participants underwent test stimulations, gradually increasing to 1 mA, to ensure comfort and tolerance without discomfort during the stimulation process.

### Participant Blinding

The study was single-blind, with all participants blinded to TI On/Off blocks. Subjects were debriefed at the end of the study and asked if they could determine when TI stimulation was active as opposed to sham. While participants reported being aware of initial TI stimulation (i.e., the first 2-3 minutes of active TI), they reported being unable to determine when TI or sham stimulation was being delivered for the remainder of the experimental session.

### Image Acquisition and Analysis

Imaging data was acquired on a Siemens 3-Tesla PRISMA scanner (Siemens AG, Munich, Germany) using a 32-channel head coil using multiband structural and functional imaging. Structural images were scanned using a single-shot, high-resolution MPRAGE sequence (TR/TE=1900/2.27ms; flip angle=9; FoV=250×250mm; 192×1.0mm slices) and functional images was acquired with T2*-weighted EPI sequences with a multiband acceleration factor of 4 (TR/TE=1000/30ms; flip angle=65; FoV=220×220mm; 52×3mm slices). Neuroimaging data was preprocessed and analyzed within a custom pipeline implementing various packages (e.g., SPM12, FSL, ICA-AROMA).

### fMRI analysis

Neuroimaging data was analyzed using standard preprocessing routines involving realignment/motion-correction, co-registration, and smoothing with a 6mm Gaussian kernel. Images were processed in native space for the purposes of TI targeting and then re-processed and normalized to MNI space using a non-linear registration technique for the purposes of group level analysis ^53^.

#### Univariate Analysis

First level contrasts were calculated using the general linear model after correcting the data with a canonical hemodynamic response. First level t-contrast maps from Color Word Stroop task were utilized to aid targeting procedure (Incongruent > Congruent trials for the dmPFC target; Congruent > Incongruent trials for the rmPFC target).

### Individualized head model

Tetrahedral head mesh model of participants were reconstructed from T1– and T2-weighted structural MRI using the “charm” pipeline from the SimNIBS framework ^54^ and segmented with iSEG by structure (e.g., skin, spongy bone, compact bone, cerebrospinal fluid, blood, gray matter, white matter).

### Electromagnetic field computation

Electromagnetic field computations were generated using the Sim4Life platform to simulate TI stimulation, which utilized ohmic-current-dominated electro-quasistatic finite-element-method solver to solve Maxwell’s equation, a four partial differential equation that describes electromagnetic fields. Subject-specific head models utilized in the simulation were generated from T1– and T2-weighted structural images and segmented with CHARM from the SimNIBS package and iSEG by structure (e.g., skin, spongy bone, compact bone, cerebrospinal fluid, blood, gray matter, white matter). Segmented images were imported and modeled within the Sim4Life environment. Two electromagnetic physics solvers were generated to simulate TI envelope. Dirichlet boundary conditions with a 1 V peak-to-peak constant potential were applied to two pairs of modeled electrodes within each simulation. Tissue-specific electrical conductivity values and density properties were applied to each segment to ensure accurate electrophysiology and 0.5 x 0.5 x 0.5 voxels were generated for a high-resolution simulation. Once both fields were calculated, maximum modulation of the two calculated fields were computed. fMRI activation maps were imported and overlayed to guide electrode placement and to verify electric field strength is maximal at target regions (rmPFC or dmPFC).

### Electrode Placement and TI Targeting

Precise participant head registration and electrode placement were achieved using the Polaris Vicra 3D tracking system (NDI, Ontario, Canada) and Brainsight software (Rogue Research, Quebec, Canada). Participants wore a head tracker, allowing the Polaris Vicra cameras to calculate head position and orientation in real-time. Brainsight software was integrated with the Polaris Vicra system, enabling real-time visualization and manipulation of 3D anatomical data. Participant MRI scans were imported into Brainsight and overlaid with the real-time tracked head position. Precise head registration was achieved by manually matching points on the participant’s head to corresponding points on their MRI scan within Brainsight software. Strategic landmarks such as nose bridge, nose tip, and right and left ear were inputted in the Brainsight software. A stylus equipped with reflective markers was then used to point at these specific landmarks on the live camera feed. This process established precise head registration within the software, ensuring accurate alignment of the participant’s head with their digital anatomical data. Pre-calculated electrode coordinates were then overlaid onto the MRI image within Brainsight, providing real-time visual guidance for electrode placement, the same stylus used above was used to ensure proper guidance. This provided accurate targeting of specific brain regions.

### Behavioral Analysis

Linear mixed effect models (LMMs) were employed to predict response times with the lme4 package in R (version 4.2.2), which utilizes the REML (Restricted Maximum Likelihood) model fit method as it provides unbiased estimates of fixed effects and better handles complex covariance structure. To ensure convergence and efficient optimization of the model parameters, the BOBYQA (Bound Optimization BY Quadratic Approximations) algorithm with a set maximum of 200,000 iterations was employed. To assess the significance of the fixed effects, t-tests were conducted using Satterthwaite’s method, which adjusts degrees of freedom to account for the complexity of the model. Statistics presented utilized the dmPFC stimulation site for Color-Word Stroop and neutral valence images for Affective Stroop as the reference factor, with releveling conducted to find additional associations (e.g., unguided dmPFC vs. rmPFC, aversive vs. pleasant stimuli).

### Analytic Strategy

Using the LLM models described above, we conducted a series of analyses were performed for each task to assess for the presence of main effects of stimulation time on RT or accuracy, as well as interactions with trial condition (incongruent or congruent), distractor valence (Affective Stroop only) and target site (rmPFC, guided dmPFC, unguided dmPFC). For each analysis, we initially performed a full-factorial model that included all main effects and higher-order interactions between TI stimulation, target site, and task conditions. Significant omnibus interactions identified by each model were then decomposed in separate models. To make pair-wise comparisons between sites, we also performed “re-leveled” models that differed only in terms of which site was arbitrarily assigned as a reference. Corrections for multiple-comparisons was achieved using a the Benjamini-Hochberg procedure to control the false discovery rate (FDR) for the total number of omnibus models performed (not including models run to decompose significant interactions). This included all main omnibus models assessing effects on RT and accuracy for block and stimulation effects for Color-Word Stroop and Affective Stroop, respectively.

## Acknowledgements

We are grateful for the help of Shosuke Suzuki, Michael Shi, and Tatiana Pillsbury in completing these studies. This work was supported by funding from the European Union’s Horizon Europe research and innovation programme under grant agreement No. 101101040 (TREATMENT) and No. 101088623 (EMUNITI) to AW and well as I3 Innovation award from Emory University to NF and MTT.

