## Supplemental Methods for "Assessing the distinct contributions of rostral and dorsomedial prefrontal cortex to cognitive control using temporal interference brain stimulation"

Temporal interference brain stimulation to the dorsomedial prefrontal cortex alters Stroop effect

**SUPPLEMENTAL METHODS**

**Participant Demographics**

**Demographics**

| **Age** |  |
| --- | --- |
| Age (Average) | 24.17647 |
| Age (Mean) | 22 |
| Age (Range) | 18-37 |
| **Sex** |  |
| Male | 42.86% |
| Female | 57.14% |
| **Race** |  |
| White | 25.71% |
| Black | 11.43% |
| Asian | 48.57% |
| Other | 8.57% |
| Undisclosed Race | 5.71% |
| **Ethnicity** |  |
| Hispanic | 5.71% |
| Non-Hispanic | 94.29% |

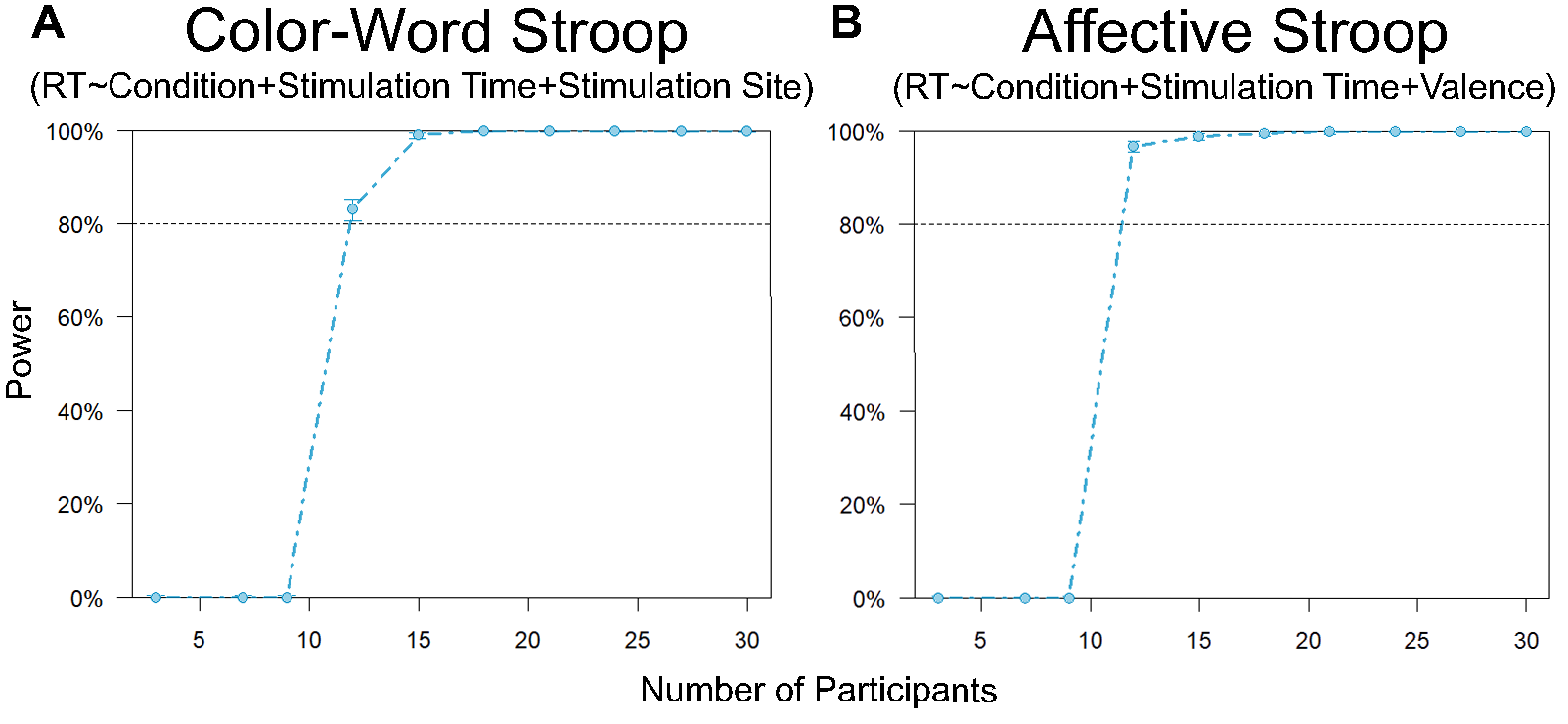

**Figure S1: Power Analysis.** Results from power analysis for **A.** Color-Word Stroop and **B.** Affective Stroop.

Power Analysis: Post-hoc power analysis conducted using the simr R package^1^ on the Color-Word Stroop and Affective Stroop data showed a power of 100%. This was determined based on 1000 simulations and utilizing the Kenward Roger test from pbkrtest^2^, using an alpha level of 0.05 and a sample size of 9557 trials for Color-Word Stroop and 5785 trials for Affective Stroop (see **Fig. S1**) and using the following simplified models:

| Task | Model |
| --- | --- |
| Color-Word Stroop | RT ~ Condition + Stimulation Time + Stimulation Site |
| Affective Stroop | RT ~ Condition + Stimulation Time + Valence |

Adverse Events: Two participants out of the 34 who received TI stimulation reported an adverse event (5.88%). One participant reported the experience of nausea during the TI stimulation session, while a second reported the experience of anxiety. In both cases, it was not clear whether the adverse event was directly related to study participation or external factors. Both events were deemed mild in severity and resolved shortly after the end of the study visit.

**SUPPLEMENTAL RESULTS**

**Neuroimaging Results: Contrast Maps**

**Color-Word Stroop: Incongruent – Congruent Trials**

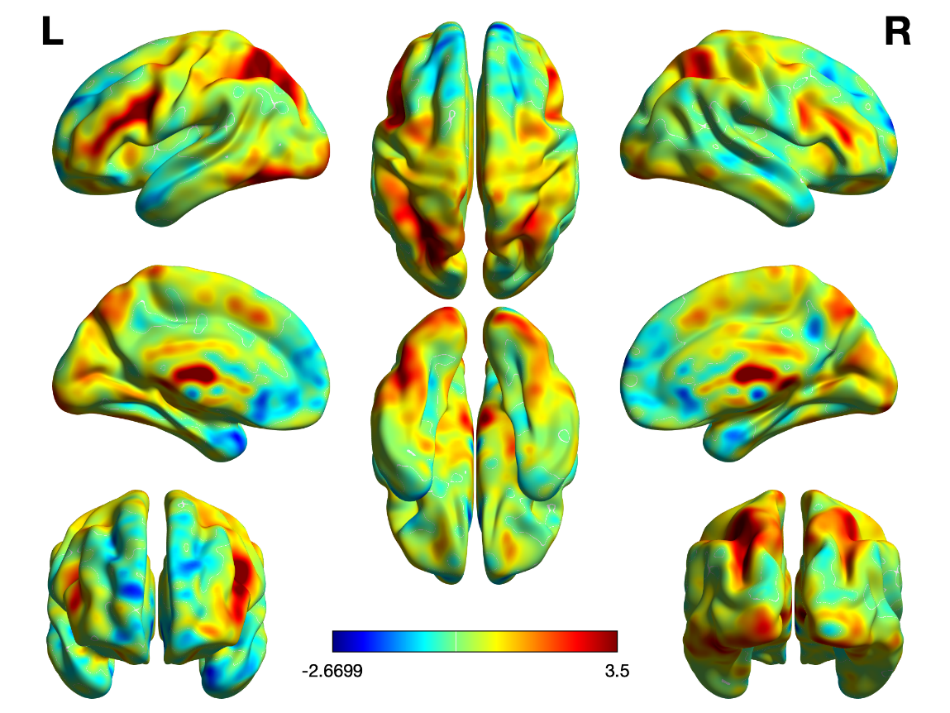

**Affective Stroop: Incongruent – Congruent Trials**

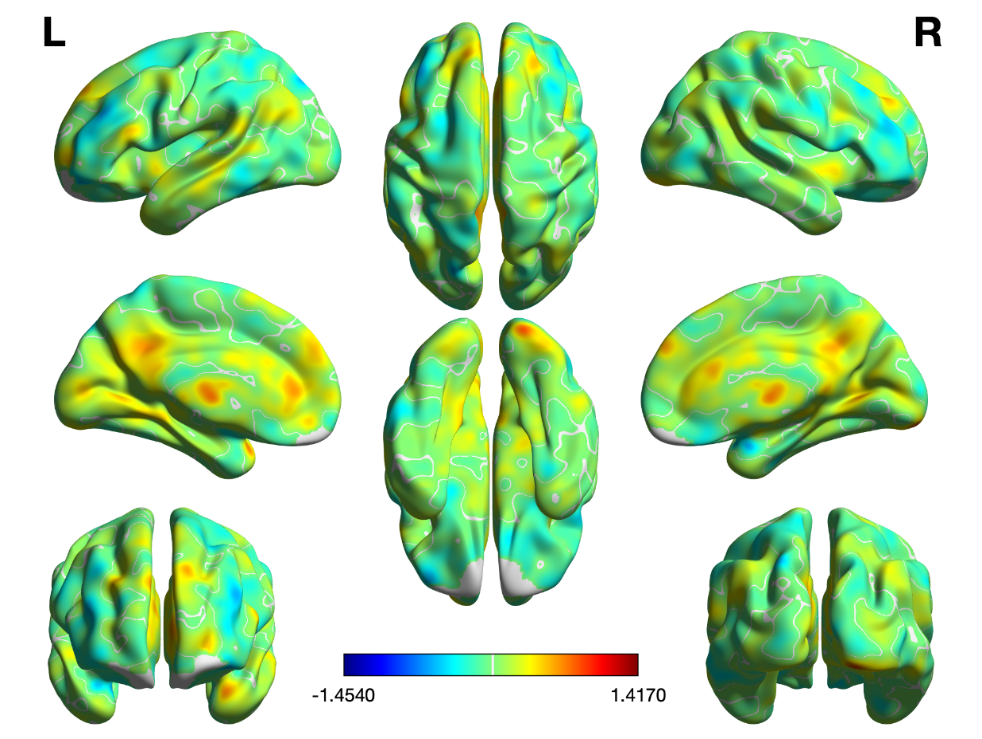

**Additional Color-Word Stroop Block Analyses**

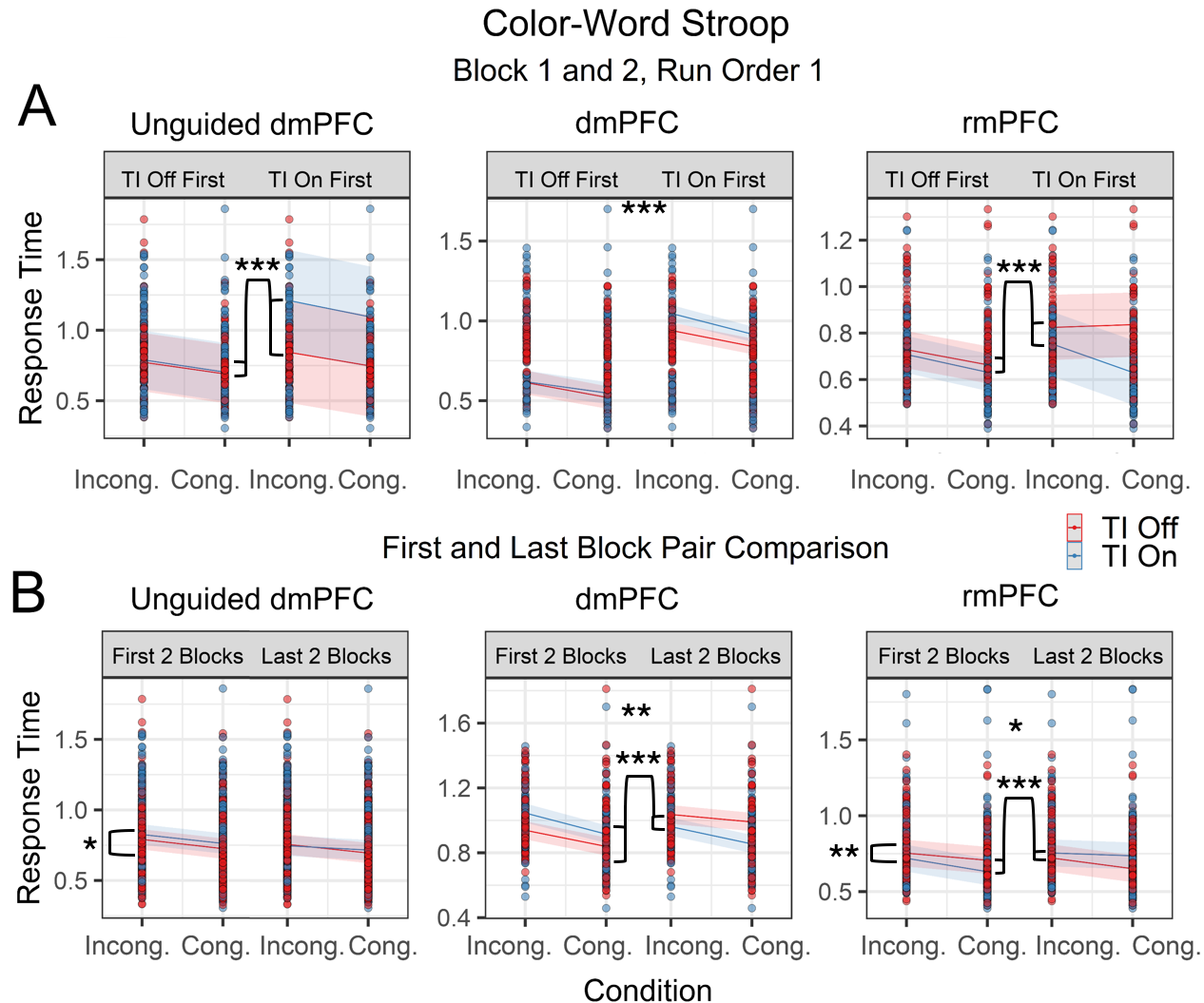

**Figure S2. Paired Block Analyses for Color-Word Stroop.** **A.** Effects of TI on response time during the first two blocks of Color-Word Stroop in individuals who ran Color-Word Stroop first, split by condition, stimulation site and stimulation order. **B.** Effects of TI on response time during the first two blocks compared to the last 2 blocks of Color-Word Stroop in individuals who ran Color-Word Stroop first. Significance codes same as below (see: Linear Mixed-Effects Models).

Additional Color-Word Stroop block analyses were conducted to further examine possible carryover effects between TI on and TI off blocks. A model conducted on blocks 1 and 2 of participants who completed Color-Word Stroop first and including stimulation order (**Fig. S2A**) revealed effects of condition (t = 1.754, p = .0797), stimulation site (dmPFC vs. unguided dmPFC: t = 2.094, p = .0837), and three way block × stimulation site × stimulation order interactions (dmPFC vs. rmPFC: t = -2.616, p = .0091; dmPFC vs. unguided dmPFC: t = 3.2504, p = .0012; unguided dmPFC vs. rmPFC: t = -5.947, p = 3.86e-9).

To assess whether there were changes in block type effects across task, an additional model was conducted looking at first and last TI On/Off block pairs within the initial target site in participants that were administered the Color-Word Stroop first. Effects of condition (*t* = 2.631, *p* = .0086), block type (*t* = 2.033, *p* = .0422), block pair (*t* = 3.97, *p* = 7.49e-5), block type × stimulation site (dmPFC vs. rmpFC: *t* = -3.311, *p* = .0009; unguided dmPFC vs. rmPFC: *t* = -4.610, *p* = 4.32e-6), block pair x stimulation site (dmPFC vs. rmpFC: *t* = -4.436, *p* = 9.74e-6; dmPFC vs. unguided dmPFC: *t* = -3.624, *p* = .0003), and a three way block type × stimulation site × block pair (dmPFC vs. rmpFC: *t* = 5.685, *p* = 1.53e-8; unguided dmPFC vs. rmPFC: *t* = 4.996, *p* = 6.46e-7) were found, further supporting the possibility of a carryover effect.

**Linear Mixed-Effects Models**

Model formula listed in the first row of each table in Wilkinson Notation. For re-leveled models, only unique comparisons were assessed for FDR. FDR only calculated for main omnibuses discussed in results.

**Significance codes:**

| p-value | Code |
| --- | --- |
| 0.001 | *** |
| 0.01 | ** |
| 0.05 | * |
| 0.1 | . |

Rounded to nearest relevant decimal.

**Variable Key:**

| **Variable** | **Description** |
| --- | --- |
| RT | Response time (ms) |
| Condition | Congruent (1) or incongruent (0) Stroop stimulus |
| TI | TI 20 Hz Δƒ (1) or 0 Hz Δƒ (0) |
| Stimulation Site | Site of stimulation (Unguided dmPFC, rmPFC, dmPFC) |
| Valence | Emotional valence of distractor during Affective Stroop (Pleasant, Aversive, Neutral); Valence2 pools emotional distractors for only two levels (Emotional, Neutral) |
| Trials | Task trial number |
| Correct | Correct (1) or incorrect (0) trial as determined by button press |
| Run Order | Order in which task was administered (1^st^, 2^nd^, or 3^rd^) |
| Stimulation Time | Cumulative amount of stimulation received across tasks in minutes. Updates every minute and end of block. |
| Stimulation Order | Whether participant received TI every odd block (1) or even block (0). |
| Block Pair | Either first (1) or last (2) TI On/Off block pair in at a given target during a task. |

**Color-Word Stroop: Block**

(Omnibus: full factorial model testing Color-Word Stroop response time based on block type)

| RT~Condition*TI*Stimulation Site +(1\|Subject) |  |  |  |  |  |  |
| --- | --- | --- | --- | --- | --- | --- |
|  | Estimate | Std.Error | df | t | Pr(>\|t\|) | FDR |
| **(Intercept)** | **0.713601** | **0.030056** | **29.83907** | **23.74233** | **6.25E-21***** | **1.06E-19** |
| **Condition** | **0.08082** | **0.008268** | **9433.021** | **9.77442** | **1.85E-22***** | **6.29E-21** |
| TI | 0.004024 | 0.008159 | 9433.022 | 0.493222 | 0.621867141 | 0.755124 |
| Unguided dmPFC Site | 0.003086 | 0.049504 | 29.52319 | 0.062333 | 0.95071768 | 0.950718 |
| **rmPFC Site** | **0.021553** | **0.008595** | **9435.658** | **2.507632** | **0.012170987**** | **0.037619** |
| Condition:TI | -0.00228 | 0.011661 | 9433.02 | -0.19549 | 0.845012344 | 0.897826 |
| **Condition:Unguided dmPFC Site** | **-0.02257** | **0.01258** | **9433.022** | **-1.79395** | **0.072853717^.^** | **0.163119** |
| Condition:rmPFC Site | -0.01213 | 0.01215 | 9433.018 | -0.99835 | 0.31813392 | 0.540828 |
| TI:Unguided dmPFC Site | 0.009849 | 0.01249 | 9433.026 | 0.788607 | 0.43036142 | 0.609679 |
| TI:rmPFC Site | -0.01698 | 0.012029 | 9433.024 | -1.4115 | 0.15813041 | 0.28297 |
| Condition:TI:Unguided dmPFC Site | -0.0088 | 0.017768 | 9433.028 | -0.49535 | 0.620363371 | 0.755124 |
| Condition:TI:rmPFC Site | 0.006281 | 0.017161 | 9433.023 | 0.366034 | 0.714348278 | 0.783479 |

**Color-Word Stroop: Block**

(Omnibus; Relevel: Unguided dmPFC)

| RT~Condition*TI*Stimulation Site+(1\|Subject) |  |  |  |  |  |  |
| --- | --- | --- | --- | --- | --- | --- |
|  | Estimate | Std.Error | df | t | Pr(>\|t\|) | FDR |
| **(Intercept)** | **0.716687** | **0.039336** | **29.34106** | **18.21983** | **1.53E-17** | N/A |
| **Condition** | **0.058252** | **0.009481** | **9433.022** | **6.144439** | **8.35E-10** | N/A |
| TI | 0.013873 | 0.009457 | 9433.028 | 1.467066 | 0.142391569 | N/A |
| dmPFC Site | -0.00309 | 0.049504 | 29.52317 | -0.06233 | 0.950717691 | N/A |
| rmPFC Site | 0.018468 | 0.049567 | 29.67332 | 0.372579 | 0.712110135 | 0.783479 |
| Condition:TI | -0.01108 | 0.013406 | 9433.034 | -0.82658 | 0.40849684 | N/A |
| **Condition:dmPFC Site** | **0.022567** | **0.01258** | **9433.022** | **1.793947** | **0.072853717** | N/A |
| Condition:rmPFC Site | 0.010437 | 0.013005 | 9433.022 | 0.802526 | 0.422268832 | 0.609679 |
| TI:dmPFC Site | -0.00985 | 0.01249 | 9433.026 | -0.78861 | 0.430361419 | N/A |
| **TI:rmPFC Site** | **-0.02683** | **0.012944** | **9433.021** | **-2.07261** | **0.038235307** | **0.092857** |
| Condition:TI:dmPFC Site | 0.008801 | 0.017768 | 9433.028 | 0.495352 | 0.620363371 | N/A |
| Condition:TI:rmPFC Site | 0.015083 | 0.018391 | 9433.026 | 0.820126 | 0.412165131 | 0.609679 |

**Color-Word Stroop: Block 1, Run 1**

(Omnibus: full factorial model testing Color-Word Stroop response time based on first block in individuals who were administered Color-Word Stroop first).

| RT~Condition*TI*Stimulation Site+(1\|Subject) |  |  |  |  |  |  |
| --- | --- | --- | --- | --- | --- | --- |
|  | Estimate | Std.Error | df | t | Pr(>\|t\|) | FDR |
| **(Intercept)** | **0.519499** | **0.123808** | **5.462165** | **4.196016** | **0.007025**** | **0.1124** |
| **Condition** | **0.093236** | **0.055405** | **463.001** | **1.682821** | **0.093084^.^** | **0.297869** |
| **TI** | **0.39551** | **0.151535** | **5.448048** | **2.610031** | **0.043909*** | **0.297869** |
| Unguided dmPFC Site | 0.166295 | 0.143222 | 5.502032 | 1.1611 | 0.293486 | 0.782629 |
| rmPFC Site | 0.143536 | 0.142914 | 5.45505 | 1.004348 | 0.35766 | 0.817509 |
| Condition:TI | 0.038364 | 0.067528 | 463.0579 | 0.56812 | 0.570229 | 0.960063 |
| Condition:Unguided dmPFC Site | -0.01305 | 0.064608 | 463.0088 | -0.20194 | 0.840055 | 0.960063 |
| Condition:rmPFC Site | -0.02776 | 0.06331 | 463.0021 | -0.43854 | 0.661201 | 0.960063 |
| TI:Unguided dmPFC Site | 0.012893 | 0.208663 | 5.489764 | 0.061787 | 0.95292 | 0.976827 |
| **TI:rmPFC Site** | **-0.42916** | **0.208296** | **5.451343** | **-2.06034** | **0.089735^.^** | **0.297869** |
| Condition:TI:Unguided dmPFC Site | -0.00271 | 0.093386 | 463.0345 | -0.02906 | 0.976827 | 0.976827 |
| Condition:TI:rmPFC Site | 0.019061 | 0.0914 | 463.0326 | 0.208538 | 0.8349 | 0.960063 |

**Color-Word Stroop: Block 1, Run 1**

(Omnibus; Relevel: Unguided dmPFC)

| RT~Condition*TI*Stimulation Site+(1\|Subject) |  |  |  |  |  |  |
| --- | --- | --- | --- | --- | --- | --- |
|  | Estimate | Std.Error | df | t | Pr(>\|t\|) | FDR |
| **(Intercept)** | **0.685794** | **0.072001** | **5.622488** | **9.524764** | **0.000112***** | N/A |
| **Condition** | **0.08019** | **0.033234** | **463.0303** | **2.412884** | **0.016214*** | N/A |
| **TI** | **0.408403** | **0.143448** | **5.536882** | **2.84704** | **0.032032*** | N/A |
| dmPFC Site | -0.16629 | 0.143222 | 5.502032 | -1.1611 | 0.293486 | N/A |
| rmPFC Site | -0.02276 | 0.101392 | 5.527716 | -0.22447 | 0.830457 | 0.960063 |
| Condition:TI | 0.03565 | 0.064505 | 463.0088 | 0.552669 | 0.580757 | N/A |
| Condition:dmPFC Site | 0.013047 | 0.064608 | 463.0088 | 0.201937 | 0.840055 | N/A |
| Condition:rmPFC Site | -0.01472 | 0.0452 | 463.019 | -0.32561 | 0.744869 | 0.960063 |
| TI:dmPFC Site | -0.01289 | 0.208663 | 5.489764 | -0.06179 | 0.95292 | N/A |
| **TI:rmPFC Site** | **-0.44205** | **0.202489** | **5.495891** | **-2.1831** | **0.075894^.^** | **0.297869** |
| Condition:TI:dmPFC Site | 0.002714 | 0.093386 | 463.0345 | 0.029062 | 0.976827 | N/A |
| Condition:TI:rmPFC Site | 0.021775 | 0.08919 | 463.0057 | 0.244135 | 0.807234 | 0.960063 |

**Color-Word Stroop: Block 1, Run 1 – dmPFC**

(Deconstruction of Color-Word Stroop: Block 1, Run 1)

| RT~Congruent*TI+(1\|Sub) |  |  |  |  |  |
| --- | --- | --- | --- | --- | --- |
|  | Estimate | Std.Error | df | t | Pr(>\|t\|) |
| **(Intercept)** | **0.519499** | **0.037715** | **128** | **13.77442** | **4.86E-27***** |
| Condition | 0.093236 | 0.055301 | 128 | 1.685986 | 0.094235 |
| **TI** | **0.39551** | **0.045869** | **128** | **8.62259** | **2.14E-14***** |
| Condition:TI | 0.036948 | 0.067362 | 128 | 0.548503 | 0.584302 |

**Color-Word Stroop: Block 1, Run 1 – rmPFC**

(Deconstruction of Color-Word Stroop: Block 1, Run 1)

| RT~Congruent*TI+(1\|Sub) |  |  |  |  |  |
| --- | --- | --- | --- | --- | --- |
|  | Estimate | Std.Error | df | t | Pr(>\|t\|) |
| **(Intercept)** | **0.662983** | **0.041249** | **2.422915** | **16.07255** | **0.001606***** |
| **Condition** | **0.065414** | **0.025333** | **180.0037** | **2.58216** | **0.010613**** |
| TI | -0.0336 | 0.082664 | 2.442402 | -0.40646 | 0.717323 |
| Condition:TI | 0.057483 | 0.050935 | 180.0007 | 1.128559 | 0.260586 |

**Color-Word Stroop: Block 1, Run 1 – unguided dmPFC**

(Deconstruction of Color-Word Stroop: Block 1, Run 1)

| RT~Congruent*TI+(1\|Sub) |  |  |  |  |  |
| --- | --- | --- | --- | --- | --- |
|  | Estimate | Std.Error | df | t | Pr(>\|t\|) |
| **(Intercept)** | **0.685648** | **0.105138** | **2.15262** | **6.521435** | **0.018809*** |
| **Condition** | **0.080281** | **0.038815** | **156.0176** | **2.068292** | **0.040263*** |
| TI | 0.408549 | 0.209757 | 2.131555 | 1.947725 | 0.18289 |
| Condition:TI | 0.035559 | 0.075337 | 156.013 | 0.471994 | 0.63759 |

**Color-Word Stroop: Block 1 and 2, Run 1**

(Omnibus: full factorial model testing Color-Word Stroop response time based on first two blocks in individuals who were administered Color-Word Stroop first).

| RT~Condition*TI*CongSite+(1\|Sub) |  |  |  |  |  |  |
| --- | --- | --- | --- | --- | --- | --- |
|  | Estimate | Std.Error | df | t | Pr(>\|t\|) | FDR |
| **(Intercept)** | **0.732242** | **0.094498** | **8.64248** | **7.748778** | **3.58E-05***** | **0.000286** |
| **Condition** | **0.096967** | **0.030955** | **945.009** | **3.132522** | **0.001786**** | **0.007144** |
| **TI** | **0.060247** | **0.030162** | **945.0039** | **1.997466** | **0.046061*** | **0.147395** |
| rmPFC Site | -0.0255 | 0.124984 | 8.635762 | -0.20401 | 0.843071 | 0.96351 |
| Unguided dmPFC Site | -0.02748 | 0.125243 | 8.707378 | -0.21938 | 0.83141 | 0.96351 |
| Condition:TI | 0.011978 | 0.04374 | 945.0349 | 0.273836 | 0.784271 | 0.96351 |
| Condition:rmPFC Site | -0.05154 | 0.040707 | 945.0094 | -1.26611 | 0.205784 | 0.548757 |
| Condition:Unguided dmPFC Site | -0.01356 | 0.04202 | 945.0104 | -0.32259 | 0.747076 | 0.96351 |
| **TI:rmPFC Site** | **-0.13621** | **0.039962** | **945.0065** | **-3.40853** | **0.000681***** | **0.003632** |
| TI:Unguided dmPFC Site | 0.039526 | 0.041383 | 945.0147 | 0.955122 | 0.33976 | 0.67952 |
| Condition:TI:rmPFC Site | 0.030622 | 0.057438 | 945.0234 | 0.53313 | 0.594069 | 0.95051 |
| Condition:TI:Unguided dmPFC Site | -0.00294 | 0.059358 | 945.0275 | -0.04955 | 0.96049 | 0.986778 |

**Color-Word Stroop: Block 1 and 2, Run 1**

(Omnibus; Relevel: Unguided dmPFC)

| RT~Condition*TI*CongSite+(1\|Sub) | | | |  |  |  |
| --- | --- | --- | --- | --- | --- | --- |
|  | Estimate | Std.Error | df | t | Pr(>\|t\|) | FDR |
| **(Intercept)** | **0.704766** | **0.082195** | **8.794287** | **8.574353** | **1.47E-05***** | N/A |
| **Condition** | **0.083412** | **0.028416** | **945.0121** | **2.935444** | **0.003411**** | N/A |
| **TI** | **0.099773** | **0.028334** | **945.027** | **3.521371** | **0.00045***** | N/A |
| dmPFC Site | 0.027476 | 0.125243 | 8.707384 | 0.21938 | 0.83141 | N/A |
| rmPFC Site | 0.001978 | 0.115962 | 8.710352 | 0.017053 | 0.986778 | 0.986778 |
| Condition:TI | 0.009036 | 0.040127 | 945.0187 | 0.225191 | 0.821879 | N/A |
| Condition:dmPFC Site | 0.013555 | 0.04202 | 945.0104 | 0.322592 | 0.747076 | N/A |
| Condition:rmPFC Site | -0.03798 | 0.038811 | 945.0111 | -0.9787 | 0.327976 | 0.67952 |
| TI:dmPFC Site | -0.03953 | 0.041383 | 945.0147 | -0.95512 | 0.33976 | N/A |
| **TI:rmPFC Site** | **-0.17574** | **0.038601** | **945.0191** | **-4.55269** | **5.99E-06***** | **9.58E-05** |
| Condition:TI:dmPFC Site | 0.002941 | 0.059358 | 945.0275 | 0.049552 | 0.96049 | N/A |
| Condition:TI:rmPFC Site | 0.033563 | 0.054737 | 945.0136 | 0.613174 | 0.539909 | 0.95051 |

**Color-Word Stroop: Block 1, Run 1 – dmPFC**

(Deconstruction of Color-Word Stroop: Block 1 and 2, Run 1)

| RT~Congruent*TI+(1\|Sub) |  |  |  |  |  |
| --- | --- | --- | --- | --- | --- |
|  | Estimate | Std.Error | df | t | Pr(>\|t\|) |
| **(Intercept)** | **0.73223** | **0.121543** | **2.087133** | **6.024458** | **0.023893*** |
| **Condition** | **0.096919** | **0.029566** | **265.0009** | **3.278028** | **0.001185***** |
| TI | 0.060259 | 0.028809 | 265.0001 | 2.0917 | 0.037417 |
| Condition:TI | 0.012091 | 0.041778 | 265.0048 | 0.289405 | 0.772498 |

**Color-Word Stroop: Block 1, Run 1 – rmPFC**

(Deconstruction of Color-Word Stroop: Block 1 and 2, Run 1)

| RT~Congruent*TI+(1\|Sub) |  |  |  |  |  |
| --- | --- | --- | --- | --- | --- |
|  | Estimate | Std.Error | df | t | Pr(>\|t\|) |
| **(Intercept)** | **0.706718** | **0.035953** | **4.159188** | **19.65652** | **2.92E-05***** |
| **Condition** | **0.045325** | **0.02317** | **366.015** | **1.956183** | **0.051204*** |
| **TI** | **-0.07593** | **0.022977** | **366.0148** | **-3.30474** | **0.001045***** |
| Condition:TI | 0.042668 | 0.03263 | 366.0109 | 1.307615 | 0.191825 |

**Color-Word Stroop: Block 1, Run 1 – unguided dmPFC**

(Deconstruction of Color-Word Stroop: Block 1 and 2, Run 1)

| RT~Congruent*TI+(1\|Sub) |  |  |  |  |  |
| --- | --- | --- | --- | --- | --- |
|  | Estimate | Std.Error | df | t | Pr(>\|t\|) |
| **(Intercept)** | **0.704765** | **0.096324** | **3.288317** | **7.316644** | **0.003839**** |
| **Condition** | **0.083413** | **0.032944** | **314.0097** | **2.531929** | **0.011831*** |
| **TI** | **0.099772** | **0.032849** | **314.0146** | **3.037266** | **0.002588**** |
| Condition:TI | 0.009036 | 0.046523 | 314.0119 | 0.194223 | 0.846127 |

**Color-Word Stroop: Block 1 and 2, Run 1 with Stimulation Order**

(Alternative Omnibus: full factorial model testing Color-Word Stroop response time based on first two blocks in individuals who were administered Color-Word Stroop first with the addition of stimulation time).

| RT~Condition*TI*Stimulation Order*CongSite+(1\|Sub) |  |  |  |  |  |
| --- | --- | --- | --- | --- | --- |
|  | Estimate | Std.Error | df | t | Pr(>\|t\|) |
| **(Intercept)** | **0.519499** | **0.124543** | **5.686944** | **4.171254** | **0.0066**** |
| **Condition** | **0.093236** | **0.053151** | **936.0034** | **1.75416** | **0.07973^.^** |
| TI | 0.027949 | 0.050727 | 936.0034 | 0.550969 | 0.581786 |
| StimOrder | 0.14365 | 0.143767 | 5.680185 | 0.999186 | 0.358349 |
| rmPFC | 0.171053 | 0.144043 | 5.723982 | 1.187513 | 0.281956 |
| **Unguided dmPFC** | **0.319356** | **0.152487** | **5.68015** | **2.094309** | **0.083728^.^** |
| Condition:TI | -0.02299 | 0.073473 | 936.0034 | -0.31297 | 0.754373 |
| Condition:Stimulation Order | -0.02794 | 0.060735 | 936.0039 | -0.46002 | 0.645606 |
| TI:Stimulation Order | -0.0115 | 0.061978 | 936.0071 | -0.18554 | 0.852845 |
| Condition:rmPFC | -0.06012 | 0.058574 | 936.0071 | -1.02647 | 0.304935 |
| Condition:Unguided dmPFC | -0.01521 | 0.059914 | 936.0124 | -0.2538 | 0.799705 |
| TI:rmPFC | 0.006478 | 0.06434 | 936.0034 | 0.100679 | 0.919827 |
| TI:Unguided dmPFC | 0.048206 | 0.062016 | 936.0042 | 0.777313 | 0.437171 |
| Stimulation Order:rmPFC | -0.14603 | 0.209443 | 5.666019 | -0.69721 | 0.513244 |
| Stimulation Order:Unguided dmPFC | -0.26276 | 0.210233 | 5.751922 | -1.24983 | 0.259807 |
| Condition:TI:Stimulation Order | 0.03403 | 0.084343 | 936.0052 | 0.403465 | 0.686698 |
| Condition:TI:rmPFC | 0.027379 | 0.086254 | 936.0102 | 0.317426 | 0.750991 |
| Condition:TI:Unguided dmPFC | 0.054521 | 0.089901 | 936.0169 | 0.606458 | 0.544358 |
| Condition:Stimulation Order:rmPFC | -0.08374 | 0.08776 | 936.0036 | -0.95416 | 0.340248 |
| Condition:Stimulation Order:Unguided dmPFC | 0.00814 | 0.090034 | 936.0051 | 0.090416 | 0.927976 |
| **TI:Stimulation Order:rmPFC** | **-0.22312** | **0.085305** | **936.0056** | **-2.61562** | **0.00905**** |
| **TI:Stimulation Order:Unguided dmPFC** | **0.286097** | **0.088019** | **936.008** | **3.250417** | **0.001193***** |
| Condition:TI:StimOrder:rmPFC | 0.069304 | 0.122979 | 936.0114 | 0.56354 | 0.573202 |
| Condition:TI:StimOrder:Unguided dmPFC | -0.03942 | 0.126042 | 936.0134 | -0.31276 | 0.754534 |

**Color-Word Stroop: Block 1 and 2, Run 1 with Stimulation Order**

(Alternative Omnibus; Relevel: unguided dmPFC site)

| RT~Condition*TI*Stimulation Order*CongSite+(1\|Sub) |  |  |  |  |  |
| --- | --- | --- | --- | --- | --- |
|  | Estimate | Std.Error | df | t | Pr(>\|t\|) |
| **(Intercept)** | **0.690552** | **0.072371** | **5.835807** | **9.541794** | **8.91E-05***** |
| **Condition** | **0.081737** | **0.031878** | **936.0173** | **2.564035** | **0.010501**** |
| TI | 0.012743 | 0.031883 | 936.0352 | 0.399673 | 0.689489 |
| StimOrder | 0.056599 | 0.144726 | 5.833195 | 0.391079 | 0.709622 |
| dmPFC | -0.17105 | 0.144043 | 5.723978 | -1.18751 | 0.281956 |
| rmPFC | -0.0274 | 0.101959 | 5.747535 | -0.26877 | 0.797492 |
| Condition:TI | 0.004384 | 0.045182 | 936.0281 | 0.097036 | 0.922719 |
| Condition:Stimulation Order | 0.014618 | 0.06298 | 936.007 | 0.23211 | 0.816503 |
| **TI:Stimulation Order** | **0.334303** | **0.062461** | **936.0117** | **5.352221** | **1.09E-07***** |
| Condition:dmPFC | 0.0115 | 0.061978 | 936.0071 | 0.185541 | 0.852845 |
| Condition:rmPFC | -0.01644 | 0.043358 | 936.0119 | -0.37917 | 0.704646 |
| TI:dmPFC | 0.015206 | 0.059914 | 936.0124 | 0.253801 | 0.799705 |
| TI:rmPFC | -0.04492 | 0.043292 | 936.0275 | -1.03756 | 0.299741 |
| Stimulation Order:dmPFC | 0.262756 | 0.210233 | 5.751918 | 1.249833 | 0.259808 |
| Stimulation Order:rmPFC | 0.11673 | 0.203862 | 5.741305 | 0.572594 | 0.588622 |
| Condition:TI:Stimulation Order | 0.0151 | 0.088343 | 936.0098 | 0.170928 | 0.864317 |
| Condition:TI:dmPFC | -0.02738 | 0.086254 | 936.0102 | -0.31743 | 0.750991 |
| Condition:TI:rmPFC | 0.00665 | 0.061293 | 936.0202 | 0.108501 | 0.913621 |
| Condition:Stimulation Order:dmPFC | -0.00814 | 0.090034 | 936.0051 | -0.09042 | 0.927976 |
| Condition:Stimulation Order:rmPFC | -0.09188 | 0.086767 | 936.0055 | -1.0589 | 0.289919 |
| **TI:Stimulation Order:dmPFC** | **-0.2861** | **0.088019** | **936.008** | **-3.25042** | **0.001193***** |
| **TI:Stimulation Order:rmPFC** | **-0.50922** | **0.085629** | **936.0095** | **-5.94686** | **3.86E-09***** |
| Condition:TI:StimOrder:dmPFC | 0.039421 | 0.126042 | 936.0134 | 0.312758 | 0.754534 |
| Condition:TI:StimOrder:rmPFC | 0.108725 | 0.121845 | 936.0076 | 0.892317 | 0.372452 |

**Color-Word Stroop: Block 1 and 2, Run 1 with Stimulation Order – dmPFC**

(Deconstruction: Color-Word Stroop: Block 1 and 2, Run 1 with Stimulation Order)

| RT~Condition*TI*Stimulation Order+(1\|Sub) |  |  |  |  |  |
| --- | --- | --- | --- | --- | --- |
|  | Estimate | Std.Error | df | t | Pr(>\|t\|) |
| **(Intercept)** | **0.519499** | **0.035697** | **263** | **14.55301** | **1.35E-35***** |
| **Condition** | **0.093236** | **0.052342** | **263** | **1.781284** | **0.07602^.^** |
| TI | 0.027949 | 0.049955 | 263 | 0.559489 | 0.576304 |
| **Stimulation Order1** | **0.31954** | **0.043564** | **263** | **7.334876** | **2.75E-12***** |
| Condition:TI | -0.02299 | 0.072354 | 263 | -0.31781 | 0.750881 |
| Condition:Stimulation Order1 | 0.006486 | 0.063361 | 263 | 0.102365 | 0.918545 |
| TI:Stimulation Order1 | 0.048021 | 0.061071 | 263 | 0.786325 | 0.432385 |
| Condition:TI:Stimulation Order1 | 0.053457 | 0.088518 | 263 | 0.603915 | 0.546421 |

**Color-Word Stroop: Block 1 and 2, Run 1 with Stimulation Order – Unguided dmPFC**

(Deconstruction: Color-Word Stroop: Block 1 and 2, Run 1 with Stimulation Order)

| RT~Condition*TI*Stimulation Order+(1\|Sub) |  |  |  |  |  |
| --- | --- | --- | --- | --- | --- |
|  | Estimate | Std.Error | df | t | Pr(>\|t\|) |
| **(Intercept)** | **0.690538** | **0.105753** | **2.186745** | **6.52971** | **0.018*** |
| **Condition** | **0.081803** | **0.035736** | **311.0089** | **2.289055** | **0.022746*** |
| TI | 0.012783 | 0.035742 | 311.0123 | 0.357655 | 0.720844 |
| Stimulation Order1 | 0.056613 | 0.211492 | 2.186153 | 0.267685 | 0.812109 |
| Condition:TI | 0.004281 | 0.05065 | 311.011 | 0.084524 | 0.932694 |
| Condition:Stimulation Order1 | 0.014552 | 0.070602 | 311.0069 | 0.206114 | 0.836837 |
| **TI:Stimulation Order1** | **0.334262** | **0.07002** | **311.0078** | **4.773785** | **2.79E-06***** |
| Condition:TI:Stimulation Order1 | 0.015203 | 0.099035 | 311.0075 | 0.153515 | 0.878092 |

**Color-Word Stroop: Block 1 and 2, Run 1 with Stimulation Order – rmPFC**

(Deconstruction: Color-Word Stroop: Block 1 and 2, Run 1 with Stimulation Order)

| RT~Condition*TI*Stimulation Order+(1\|Sub) |  |  |  |  |  |
| --- | --- | --- | --- | --- | --- |
|  | Estimate | Std.Error | df | t | Pr(>\|t\|) |
| **(Intercept)** | **0.66311** | **0.040817** | **2.787921** | **16.24582** | **0.000761***** |
| **Condition** | **0.065264** | **0.026294** | **363.0027** | **2.482075** | **0.013514**** |
| TI | -0.03212 | 0.026202 | 363.0163 | -1.22585 | 0.221049 |
| Stimulation Order1 | 0.173368 | 0.081547 | 2.775995 | 2.125993 | 0.130744 |
| Condition:TI | 0.011051 | 0.037056 | 363.0082 | 0.298208 | 0.765715 |
| Condition:Stimulation Order1 | -0.07723 | 0.053398 | 363.0009 | -1.44624 | 0.148974 |
| **TI:Stimulation Order1** | **-0.17497** | **0.052406** | **363.0043** | **-3.3388** | **0.000929***** |
| **Condition:TI:Stimulation Order1** | **0.123809** | **0.075079** | **363.0023** | **1.649047** | **0.100003.** |

**Color-Word Stroop: First and Last TI On/Off Pairs, Run 1**

(Supplemental Omnibus: full factorial model testing Color-Word Stroop response time comparing first and last TI On/Off block pairs in individuals who were administered Color-Word Stroop first with the addition of stimulation time)

| RT~Congruent*TI*Stimulation Site*BlockPair+(1\|Sub) | | | | | |
| --- | --- | --- | --- | --- | --- |
|  | Estimate | Std.Error | df | t | Pr(>\|t\|) |
| **(Intercept)** | **0.839185** | **0.085334** | **8.326981** | **9.834082** | **7.24E-06***** |
| **Condition** | **0.099729** | **0.037908** | **1715.996** | **2.630795** | **0.008595**** |
| **TI** | **0.075825** | **0.037299** | **1715.997** | **2.032912** | **0.042215*** |
| rmPFC | -0.13272 | 0.104495 | 8.321336 | -1.27013 | 0.238403 |
| unguided dmPFC | -0.13019 | 0.104809 | 8.421626 | -1.24212 | 0.247666 |
| **BlockPair** | **0.150522** | **0.037917** | **1716.019** | **3.969724** | **7.49E-05***** |
| Condition:TI | 0.029621 | 0.054152 | 1716.018 | 0.546995 | 0.584453 |
| Condition:rmPFC | -0.05329 | 0.046389 | 1715.998 | -1.14873 | 0.250829 |
| Condition:unguided dmPFC | -0.01441 | 0.047572 | 1715.998 | -0.303 | 0.761928 |
| **TI:rmPFC** | **-0.15154** | **0.045763** | **1715.999** | **-3.31136** | **0.000948***** |
| TI:unguided dmPFC | 0.028445 | 0.047035 | 1716.003 | 0.60477 | 0.545412 |
| Condition:BlockPair | -0.0541 | 0.054219 | 1715.999 | -0.99774 | 0.318547 |
| **TI:BlockPair** | **-0.21121** | **0.053326** | **1715.999** | **-3.96067** | **7.78E-05***** |
| **rmPFC:BlockPair** | **-0.20624** | **0.046489** | **1716.019** | **-4.43622** | **9.74E-06***** |
| **unguided dmPFC:BlockPair** | **-0.17137** | **0.04728** | **1716.024** | **-3.62451** | **0.000298***** |
| Condition:TI:rmPFC | 0.012503 | 0.065957 | 1716.012 | 0.189563 | 0.849674 |
| Condition:TI:unguided dmPFC | -0.02459 | 0.067672 | 1716.014 | -0.36339 | 0.716358 |
| Condition:TI:BlockPair | 0.030915 | 0.076632 | 1716.018 | 0.403417 | 0.686692 |
| Condition:rmPFC:BlockPair | 0.077936 | 0.066583 | 1716.004 | 1.170513 | 0.241957 |
| Condition:unguided dmPFC:BlockPair | 0.050657 | 0.067334 | 1715.999 | 0.752326 | 0.451958 |
| **TI:rmPFC:BlockPair** | **0.371653** | **0.065374** | **1716.001** | **5.685012** | **1.53E-08***** |
| TI:unguided dmPFC:BlockPair | 0.096927 | 0.066615 | 1716.001 | 1.455046 | 0.145839 |
| Condition:TI:rmPFC:BlockPair | -0.12445 | 0.093812 | 1716.016 | -1.32659 | 0.18482 |
| Condition:TI:unguided dmPFC:BlockPair | -0.10799 | 0.095234 | 1716.011 | -1.13393 | 0.256983 |

**Color-Word Stroop: First and Last TI On/Off Pairs, Run 1**

(Supplemental Omnibus; Relevel: unguided dmPFC)

| RT~Congruent*TI*CongSite*BlockPair+(1\|Sub) | | | | | |
| --- | --- | --- | --- | --- | --- |
|  | Estimate | Std.Error | df | t | Pr(>\|t\|) |
| **(Intercept)** | **0.708999** | **0.060852** | **8.612497** | **11.6513** | **1.45E-06***** |
| **Condition** | **0.085314** | **0.028741** | **1716.003** | **2.968427** | **0.003035**** |
| **TI** | **0.10427** | **0.028654** | **1716.013** | **3.638887** | **0.000282***** |
| dmPFC | 0.130185 | 0.104809 | 8.421613 | 1.242124 | 0.247666 |
| rmPFC | -0.00254 | 0.085675 | 8.460596 | -0.02961 | 0.977068 |
| BlockPair | -0.02084 | 0.028242 | 1716.033 | -0.73804 | 0.460591 |
| Condition:TI | 0.005029 | 0.040584 | 1716.008 | 0.123922 | 0.901391 |
| Condition:dmPFC | 0.014414 | 0.047572 | 1715.998 | 0.302998 | 0.761928 |
| Condition:rmPFC | -0.03887 | 0.039255 | 1716.003 | -0.9903 | 0.322165 |
| TI:dmPFC | -0.02845 | 0.047035 | 1716.003 | -0.60477 | 0.545412 |
| **TI:rmPFC** | **-0.17998** | **0.03904** | **1716.008** | **-4.61019** | **4.32E-06***** |
| Condition:BlockPair | -0.00344 | 0.039928 | 1715.998 | -0.08612 | 0.93138 |
| **TI:BlockPair** | **-0.11428** | **0.039923** | **1716.005** | **-2.86243** | **0.004255**** |
| **dmPFC:BlockPair** | **0.171366** | **0.04728** | **1716.024** | **3.624513** | **0.000298***** |
| rmPFC:BlockPair | -0.03487 | 0.039002 | 1716.027 | -0.89407 | 0.371409 |
| Condition:TI:dmPFC | 0.024591 | 0.067672 | 1716.014 | 0.363391 | 0.716358 |
| Condition:TI:rmPFC | 0.037094 | 0.055363 | 1716.005 | 0.670026 | 0.502932 |
| Condition:TI:BlockPair | -0.07707 | 0.056543 | 1715.999 | -1.36311 | 0.173026 |
| Condition:dmPFC:BlockPair | -0.05066 | 0.067334 | 1715.999 | -0.75233 | 0.451958 |
| Condition:rmPFC:BlockPair | 0.027279 | 0.055569 | 1716.005 | 0.490905 | 0.623556 |
| TI:dmPFC:BlockPair | -0.09693 | 0.066615 | 1716.001 | -1.45505 | 0.145839 |
| **TI:rmPFC:BlockPair** | **0.274726** | **0.054991** | **1716.006** | **4.995813** | **6.46E-07***** |
| Condition:TI:dmPFC:BlockPair | 0.107989 | 0.095234 | 1716.011 | 1.133929 | 0.256983 |
| Condition:TI:rmPFC:BlockPair | -0.01646 | 0.078265 | 1716.006 | -0.21033 | 0.833431 |

**Color-Word Stroop: First and Last TI On/Off Pairs, Run 1 – dmPFC**

(Deconstruction of: Color-Word Stroop: First and Last TI On/Off Pairs, Run 1)

| RT~Condition*TI*Block Pair+(1\|Sub) |  |  |  |  |  |
| --- | --- | --- | --- | --- | --- |
|  | Estimate | Std.Error | df | t | Pr(>\|t\|) |
| **(Intercept)** | **0.839039** | **0.028175** | **353** | **29.77941** | **2.54E-98***** |
| **Condition** | **0.099722** | **0.040286** | **353** | **2.475356** | **0.013779**** |
| **TI** | **0.07597** | **0.039638** | **353** | **1.916628** | **0.056092^.^** |
| **Block Pair2** | **0.151124** | **0.040286** | **353** | **3.751282** | **0.000206***** |
| Condition:TI | 0.030462 | 0.057535 | 353 | 0.529461 | 0.596819 |
| Condition:Block Pair2 | -0.05375 | 0.057617 | 353 | -0.93287 | 0.351523 |
| **TI:Block Pair2** | **-0.21151** | **0.056669** | **353** | **-3.73239** | **0.000221***** |
| Condition:TI:Block Pair2 | 0.029733 | 0.08142 | 353 | 0.36518 | 0.715195 |

**Color-Word Stroop: First and Last TI On/Off Pairs, Run 1 – rmPFC**

(Deconstruction of: Color-Word Stroop: First and Last TI On/Off Pairs, Run 1)

| RT~Condition*TI*Block Pair+(1\|Sub) |  |  |  |  |  |
| --- | --- | --- | --- | --- | --- |
|  | Estimate | Std.Error | df | t | Pr(>\|t\|) |
| (Intercept) | 0.706461 | 0.045012 | 3.975816 | 15.69509 | 0.0001 |
| Condition | 0.04641 | 0.025141 | 713.0025 | 1.845988 | 0.065308 |
| TI | -0.07571 | 0.024932 | 713.0022 | -3.03672 | 0.002479 |
| Block Pair2 | -0.05577 | 0.025291 | 713.0143 | -2.20504 | 0.02777 |
| Condition:TI | 0.042142 | 0.035406 | 713.0007 | 1.190257 | 0.234341 |
| Condition:Block Pair2 | 0.023841 | 0.036339 | 713.0095 | 0.656062 | 0.511996 |
| TI:Block Pair2 | 0.160487 | 0.035558 | 713.0054 | 4.513356 | 7.46E-06 |
| Condition:TI:Block Pair2 | -0.09359 | 0.050881 | 713.0094 | -1.83939 | 0.066273 |

**Color-Word Stroop: First and Last TI On/Off Pairs, Run 1 – Unguided dmPFC**

(Deconstruction of: Color-Word Stroop: First and Last TI On/Off Pairs, Run 1)

| RT~Condition*TI*Block Pair+(1\|Sub) |  |  |  |  |  |
| --- | --- | --- | --- | --- | --- |
|  | Estimate | Std.Error | df | t | Pr(>\|t\|) |
| **(Intercept)** | **0.725627** | **0.036831** | **11.933** | **19.70164** | **1.81E-10***** |
| **Condition** | **0.065308** | **0.016201** | **1822.011** | **4.031207** | **5.78E-05***** |
| **TI** | **0.036809** | **0.01613** | **1822.008** | **2.281983** | **0.022605*** |
| **Block Pair2** | **-0.02995** | **0.01615** | **1822.025** | **-1.85462** | **0.063812^.^** |
| Condition:TI | -0.00209 | 0.022875 | 1822.016 | -0.09116 | 0.927375 |
| Condition:Block Pair2 | -0.00398 | 0.022811 | 1822.003 | -0.17438 | 0.861585 |
| TI:Block Pair2 | -0.01577 | 0.022811 | 1822.009 | -0.69138 | 0.489413 |
| Condition:TI:Block Pair2 | -0.02851 | 0.032296 | 1822.008 | -0.88286 | 0.377426 |

**Affective Stroop: Block**

(Omnibus: full factorial model testing Affective Stroop response time based on block type)

| RT~Condition*TI*Valence*Stimulation Site+(1\|Subject) |  |  |  |  |  |  |
| --- | --- | --- | --- | --- | --- | --- |
|  | Estimate | Std.Error | df | t | Pr(>\|t\|) | FDR |
| **(Intercept)** | **0.562515** | **0.017002** | **35.83543** | **33.08529** | **1.86E-28***** | **9.49E-27** |
| **Condition** | **0.083075** | **0.008768** | **5636.443** | **9.474869** | **3.83E-21***** | **6.51E-20** |
| TI | 0.013786 | 0.008652 | 5636.169 | 1.593247 | 0.111161 | 0.36757 |
| Aversive | 0.009472 | 0.008773 | 5636.352 | 1.07968 | 0.280331 | 0.621604 |
| Pleasant | 0.013695 | 0.008695 | 5636.352 | 1.574983 | 0.115316 | 0.36757 |
| Unguided dmPFC Site | 0.039442 | 0.028145 | 36.17994 | 1.401384 | 0.169619 | 0.508857 |
| Condition:TI | -0.0035 | 0.012391 | 5636.251 | -0.28225 | 0.777763 | 0.922256 |
| Condition:Aversive | -0.00408 | 0.012444 | 5636.644 | -0.32771 | 0.743143 | 0.922256 |
| Condition:Pleasant | -0.00387 | 0.012412 | 5636.723 | -0.31152 | 0.755416 | 0.922256 |
| TI:Aversive | 0.002532 | 0.01246 | 5636.321 | 0.203198 | 0.838987 | 0.922256 |
| TI:Pleasant | -0.00253 | 0.01237 | 5636.289 | -0.20421 | 0.838195 | 0.922256 |
| Condition:Unguided dmPFC Site | 0.014432 | 0.014354 | 5636.582 | 1.00543 | 0.314733 | 0.668808 |
| **TI:Unguided dmPFC Site** | **-0.02657** | **0.014433** | **5636.246** | **-1.84117** | **0.065649^.^** | **0.279008** |
| Aversive:Unguided dmPFC Site | 0.002982 | 0.014406 | 5636.336 | 0.207005 | 0.836013 | 0.922256 |
| Pleasant:Unguided dmPFC Site | -0.01318 | 0.014567 | 5636.551 | -0.90477 | 0.365628 | 0.745881 |
| Condition:TI:Aversive | 0.005119 | 0.01762 | 5636.543 | 0.290528 | 0.771423 | 0.922256 |
| Condition:TI:Pleasant | -0.00229 | 0.017538 | 5636.585 | -0.13053 | 0.896154 | 0.952164 |
| Condition:TI:Unguided dmPFC Site | -0.00839 | 0.020342 | 5636.49 | -0.41263 | 0.679897 | 0.922256 |
| Condition:Aversive:Unguided dmPFC Site | -0.01082 | 0.020356 | 5636.614 | -0.53131 | 0.595224 | 0.91158 |
| Condition:Pleasant:Unguided dmPFC Site | -0.01116 | 0.020338 | 5637.09 | -0.54855 | 0.583333 | 0.91158 |
| TI:Aversive:Unguided dmPFC Site | -0.00077 | 0.020485 | 5636.435 | -0.03746 | 0.970118 | 0.986635 |
| TI:Pleasant:Unguided dmPFC Site | 0.013922 | 0.020503 | 5636.394 | 0.679028 | 0.497148 | 0.878248 |
| Condition:TI:Aversive:Unguided dmPFC Site | 0.0148 | 0.028829 | 5636.788 | 0.513359 | 0.60772 | 0.91158 |
| Condition:TI:Pleasant:Unguided dmPFC Site | 0.009354 | 0.028748 | 5636.772 | 0.325373 | 0.744911 | 0.922256 |

**Affective Stroop: Block**

(Omnibus; Relevel: Aversive Trials)

| RT~Condition*TI*Valence*Stimulation Site+(1\|Subject) | | | | |  |  |
| --- | --- | --- | --- | --- | --- | --- |
|  | Estimate | Std.Error | df | t | Pr(>\|t\|) | FDR |
| **(Intercept)** | **0.571987** | **0.0171** | **36.6714** | **33.44886** | **4.45E-29***** | N/A |
| **Condition** | **0.078998** | **0.00882** | **5636.544** | **8.956297** | **4.49E-19***** | N/A |
| **TI** | **0.016317** | **0.00896** | **5636.279** | **1.821086** | **0.068647^.^** | N/A |
| Neutral | -0.00947 | 0.008773 | 5636.352 | -1.07968 | 0.280331 | N/A |
| Pleasant | 0.004223 | 0.008887 | 5636.384 | 0.475127 | 0.634715 | 0.922256 |
| Unguided dmPFC Site | 0.042424 | 0.028131 | 36.10717 | 1.508098 | 0.14023 | N/A |
| Condition:TI | 0.001622 | 0.012517 | 5636.599 | 0.129562 | 0.896917 | N/A |
| Condition:Neutral | 0.004078 | 0.012444 | 5636.644 | 0.32771 | 0.743143 | N/A |
| Condition:Pleasant | 0.000211 | 0.012454 | 5636.878 | 0.016964 | 0.986466 | 0.986635 |
| TI:Neutral | -0.00253 | 0.01246 | 5636.321 | -0.2032 | 0.838987 | N/A |
| TI:Pleasant | -0.00506 | 0.012592 | 5636.433 | -0.40169 | 0.687928 | 0.922256 |
| Condition:Unguided dmPFC Site | 0.003617 | 0.014418 | 5636.355 | 0.250869 | 0.801925 | N/A |
| TI:Unguided dmPFC Site | -0.02734 | 0.01452 | 5636.298 | -1.883 | 0.059752 | N/A |
| Neutral:Unguided dmPFC Site | -0.00298 | 0.014406 | 5636.336 | -0.20701 | 0.836013 | N/A |
| Pleasant:Unguided dmPFC Site | -0.01616 | 0.014531 | 5636.347 | -1.11222 | 0.266093 | 0.616852 |
| Condition:TI:Neutral | -0.00512 | 0.01762 | 5636.543 | -0.29053 | 0.771423 | N/A |
| Condition:TI:Pleasant | -0.00741 | 0.017644 | 5637.009 | -0.41989 | 0.674582 | 0.922256 |
| Condition:TI:Unguided dmPFC Site | 0.006406 | 0.020388 | 5636.527 | 0.314218 | 0.753367 | N/A |
| Condition:Neutral:Unguided dmPFC Site | 0.010815 | 0.020356 | 5636.614 | 0.531311 | 0.595224 | N/A |
| Condition:Pleasant:Unguided dmPFC Site | -0.00034 | 0.020364 | 5636.716 | -0.01675 | 0.986635 | 0.986635 |
| TI:Neutral:Unguided dmPFC Site | 0.000767 | 0.020485 | 5636.435 | 0.037462 | 0.970118 | N/A |
| TI:Pleasant:Unguided dmPFC Site | 0.014689 | 0.020563 | 5636.401 | 0.714367 | 0.47503 | 0.878248 |
| Condition:TI:Neutral:Unguided dmPFC Site | -0.0148 | 0.028829 | 5636.788 | -0.51336 | 0.60772 | N/A |
| Condition:TI:Pleasant:Unguided dmPFC Site | -0.00545 | 0.028779 | 5636.777 | -0.18923 | 0.849922 | 0.922256 |

**Affective Stroop: Block 1, Run 1**

(Omnibus: full factorial model testing Affective Stroop response time based on first block in individuals who were administered Affective Stroop first)

| RT~Condition*TI*Valence*Stimulation Site+(1\|Subject) |  |  |  |  |  |  |
| --- | --- | --- | --- | --- | --- | --- |
|  | Estimate | Std.Error | df | t | Pr(>\|t\|) | FDR |
| **(Intercept)** | **0.516364** | **0.059557** | **7.326029** | **8.670052** | **4.16E-05***** | **0.001331** |
| **Condition** | **0.102029** | **0.037899** | **256.0047** | **2.692141** | **0.007568**** | **0.121088** |
| TI | 0.056685 | 0.085822 | 7.893634 | 0.66049 | 0.527739 | 0.886852 |
| Aversive | 0.04871 | 0.04007 | 256.1309 | 1.215601 | 0.225256 | 0.65529 |
| **Pleasant** | **0.062634** | **0.036944** | **256.1009** | **1.695344** | **0.091226^.^** | **0.486539** |
| Unguided dmPFC Site | 0.186204 | 0.119114 | 7.326029 | 1.563233 | 0.160074 | 0.569152 |
| Condition:TI | -0.01954 | 0.054569 | 256.0124 | -0.3581 | 0.720567 | 0.886852 |
| Condition:Aversive | -0.03364 | 0.05359 | 256.1941 | -0.62767 | 0.530777 | 0.886852 |
| Condition:Pleasant | -0.03112 | 0.05502 | 256.346 | -0.56558 | 0.572173 | 0.886852 |
| TI:Aversive | -0.02987 | 0.057293 | 256.1473 | -0.52133 | 0.602587 | 0.886852 |
| TI:Pleasant | -0.0163 | 0.054736 | 256.0673 | -0.29778 | 0.766109 | 0.88902 |
| Condition:Unguided dmPFC Site | 0.045302 | 0.076692 | 255.9947 | 0.590709 | 0.555236 | 0.886852 |
| TI:Unguided dmPFC Site | -0.21628 | 0.152125 | 7.380526 | -1.42172 | 0.195958 | 0.627066 |
| Aversive:Unguided dmPFC Site | -0.11597 | 0.075369 | 256.031 | -1.53874 | 0.125103 | 0.569152 |
| Pleasant:Unguided dmPFC Site | -0.03239 | 0.079786 | 256.015 | -0.40593 | 0.685134 | 0.886852 |
| Condition:TI:Aversive | 0.072526 | 0.076817 | 256.249 | 0.944136 | 0.345989 | 0.813696 |
| Condition:TI:Pleasant | 0.014653 | 0.077159 | 256.2791 | 0.189908 | 0.849531 | 0.906166 |
| Condition:TI:Unguided dmPFC Site | -0.02411 | 0.097909 | 256.003 | -0.24627 | 0.805674 | 0.88902 |
| Condition:Aversive:Unguided dmPFC Site | 0.050009 | 0.106733 | 256.0429 | 0.468541 | 0.639796 | 0.886852 |
| Condition:Pleasant:Unguided dmPFC Site | -0.0504 | 0.110014 | 256.0813 | -0.45813 | 0.647247 | 0.886852 |
| **TI:Aversive:Unguided dmPFC Site** | **0.192754** | **0.097661** | **256.0903** | **1.973718** | **0.049489*** | **0.31673** |
| TI:Pleasant:Unguided dmPFC Site | -0.00609 | 0.099732 | 256.0433 | -0.06103 | 0.951386 | 0.963874 |
| Condition:TI:Aversive:Unguided dmPFC Site | -0.19431 | 0.137001 | 256.1041 | -1.41833 | 0.157311 | 0.569152 |
| Condition:TI:Pleasant:Unguided dmPFC Site | 0.089845 | 0.138875 | 256.1668 | 0.64695 | 0.518244 | 0.886852 |

**Affective Stroop: Block 1, Run 1**

(Omnibus; Relevel: Pleasant Trials)

| RT~Condition*TI*Valence*Stimulation Site+(1\|Subject) | | | | | | |
| --- | --- | --- | --- | --- | --- | --- |
|  | Estimate | Std.Error | df | t | Pr(>\|t\|) | FDR |
| **(Intercept)** | **0.578998** | **0.058568** | **6.851909** | **9.885922** | **2.66E-05** | N/A |
| **Condition** | **0.070911** | **0.03981** | **256.5672** | **1.781232** | **0.076058** | N/A |
| TI | 0.040385 | 0.082841 | 6.856019 | 0.487504 | 0.641101 | N/A |
| **Neutral** | **-0.06263** | **0.036944** | **256.1009** | **-1.69534** | **0.091226** | N/A |
| Aversive | -0.01392 | 0.038783 | 256.4628 | -0.35902 | 0.719873 | 0.886852 |
| Unguided dmPFC Site | 0.153816 | 0.120942 | 7.783183 | 1.271814 | 0.240122 | N/A |
| Condition:TI | -0.00489 | 0.054641 | 256.6092 | -0.08945 | 0.928794 | N/A |
| Condition:Neutral | 0.031118 | 0.05502 | 256.346 | 0.565582 | 0.572173 | N/A |
| Condition:Aversive | -0.00252 | 0.055548 | 256.9176 | -0.04534 | 0.963874 | 0.963874 |
| TI:Neutral | 0.0163 | 0.054736 | 256.0673 | 0.297785 | 0.766109 | N/A |
| TI:Aversive | -0.01357 | 0.05295 | 256.4681 | -0.25626 | 0.797959 | 0.88902 |
| Condition:Unguided dmPFC Site | -0.0051 | 0.078837 | 256.1414 | -0.06467 | 0.94849 | N/A |
| TI:Unguided dmPFC Site | -0.22237 | 0.152617 | 7.475073 | -1.45701 | 0.1858 | N/A |
| Neutral:Unguided dmPFC Site | 0.032387 | 0.079786 | 256.015 | 0.405928 | 0.685134 | N/A |
| Aversive:Unguided dmPFC Site | -0.08359 | 0.078324 | 256.109 | -1.06718 | 0.286895 | 0.765053 |
| Condition:TI:Neutral | -0.01465 | 0.077159 | 256.2791 | -0.18991 | 0.849531 | N/A |
| Condition:TI:Aversive | 0.057872 | 0.077648 | 256.9568 | 0.745321 | 0.456759 | 0.886852 |
| Condition:TI:Unguided dmPFC Site | 0.065733 | 0.098431 | 256.3023 | 0.667808 | 0.504857 | N/A |
| Condition:Neutral:Unguided dmPFC Site | 0.050401 | 0.110014 | 256.0813 | 0.458131 | 0.647247 | N/A |
| Condition:Aversive:Unguided dmPFC Site | 0.100409 | 0.108586 | 256.2424 | 0.9247 | 0.355992 | 0.813696 |
| TI:Neutral:Unguided dmPFC Site | 0.006086 | 0.099732 | 256.0433 | 0.061027 | 0.951386 | N/A |
| **TI:Aversive:Unguided dmPFC Site** | **0.198841** | **0.098709** | **256.2744** | **2.014414** | **0.045011** | **0.31673** |
| Condition:TI:Neutral:Unguided dmPFC Site | -0.08984 | 0.138875 | 256.1668 | -0.64695 | 0.518244 | N/A |
| **Condition:TI:Aversive:Unguided dmPFC Site** | **-0.28416** | **0.138228** | **256.5198** | **-2.05571** | **0.040823** | **0.31673** |

**Affective Stroop: Block 1 and 2, Run 1**

(Omnibus: full factorial model testing affective Stroop response time based on first two blocks in individuals who were administered affective Stroop first).

| RT~Condition*TI*Valence*Stimulation Site+(1\|Sub) | | | | | | |
| --- | --- | --- | --- | --- | --- | --- |
|  | Estimate | Std.Error | df | t | Pr(>\|t\|) | FDR |
| **(Intercept)** | **0.5371** | **0.036887** | **12.96613** | **14.56079** | **2.06E-09***** | **6.59E-08** |
| **Condition** | **0.07994** | **0.027154** | **539.0054** | **2.943944** | **0.00338**** | **0.05408** |
| TI | 0.028697 | 0.028557 | 539.0697 | 1.004885 | 0.315403 | 0.776377 |
| Aversive | 0.015271 | 0.028296 | 539.0786 | 0.539702 | 0.589625 | 0.918928 |
| Pleasant | 0.03301 | 0.026639 | 539.0555 | 1.239157 | 0.215827 | 0.690646 |
| **Unguided dmPFC Site** | **0.140624** | **0.063312** | **12.50653** | **2.221121** | **0.045492*** | **0.207963** |
| Condition:TI | 0.009555 | 0.038359 | 539.1214 | 0.249092 | 0.803384 | 0.918928 |
| Condition:Aversive | 0.0101 | 0.038178 | 539.1484 | 0.264541 | 0.791464 | 0.918928 |
| Condition:Pleasant | -0.00883 | 0.03849 | 539.1532 | -0.2294 | 0.818649 | 0.918928 |
| TI:Aversive | -0.00449 | 0.03932 | 539.0144 | -0.11431 | 0.909036 | 0.928209 |
| TI:Pleasant | -0.02319 | 0.038687 | 539.2538 | -0.59954 | 0.549068 | 0.918928 |
| Condition:Unguided dmPFC Site | -0.01136 | 0.046486 | 539.0201 | -0.24445 | 0.806973 | 0.918928 |
| **TI:Unguided dmPFC Site** | **-0.12385** | **0.046467** | **539.0308** | **-2.66527** | **0.007923**** | **0.084512** |
| **Aversive:Unguided dmPFC Site** | **-0.10161** | **0.049744** | **539.2267** | **-2.04267** | **0.041571*** | **0.207963** |
| Pleasant:Unguided dmPFC Site | -0.09319 | 0.044972 | 539.0657 | -2.07209 | 0.038731 | 0.207963 |
| Condition:TI:Aversive | -0.01439 | 0.054109 | 539.0116 | -0.26595 | 0.790376 | 0.918928 |
| Condition:TI:Pleasant | -0.0049 | 0.054331 | 539.5467 | -0.09014 | 0.928209 | 0.928209 |
| Condition:TI:Unguided dmPFC Site | -0.01608 | 0.066343 | 539.0405 | -0.24233 | 0.80862 | 0.918928 |
| Condition:Aversive:Unguided dmPFC Site | 0.062239 | 0.066519 | 539.2649 | 0.93566 | 0.349867 | 0.799696 |
| Condition:Pleasant:Unguided dmPFC Site | 0.016187 | 0.066893 | 539.2345 | 0.241982 | 0.808886 | 0.918928 |
| **TI:Aversive:Unguided dmPFC Site** | **0.170238** | **0.068375** | **539.1863** | **2.489749** | **0.013084**** | **0.104672** |
| TI:Pleasant:Unguided dmPFC Site | 0.09251 | 0.064455 | 539.1276 | 1.435256 | 0.151794 | 0.539712 |
| Condition:TI:Aversive:Unguided dmPFC Site | -0.09626 | 0.094545 | 539.1559 | -1.01812 | 0.309077 | 0.776377 |
| Condition:TI:Pleasant:Unguided dmPFC Site | 0.051119 | 0.094066 | 539.3343 | 0.543435 | 0.587055 | 0.918928 |

**Affective Stroop: Block 1 and 2, Run 1**

(Omnibus; Relevel: Pleasant Trials)

| RT~Condition*TI*Valence*Stimulation Site+(1\|Sub) | | | | | | |
| --- | --- | --- | --- | --- | --- | --- |
|  | Estimate | Std.Error | df | t | Pr(>\|t\|) | FDR |
| **(Intercept)** | **0.57011** | **0.035851** | **11.57625** | **15.90237** | **3.19E-09***** | N/A |
| **Condition** | **0.071111** | **0.027273** | **539.2858** | **2.607359** | **0.009377**** | N/A |
| TI | 0.005503 | 0.02602 | 539.2696 | 0.211476 | 0.832596 | N/A |
| Neutral | -0.03301 | 0.026639 | 539.0555 | -1.23916 | 0.215827 | N/A |
| Aversive | -0.01774 | 0.026975 | 539.2383 | -0.65761 | 0.511067 | 0.918928 |
| Unguided dmPFC Site | 0.047439 | 0.061845 | 11.3909 | 0.76706 | 0.45865 | N/A |
| Condition:TI | 0.004658 | 0.038223 | 539.5317 | 0.121852 | 0.903062 | N/A |
| Condition:Neutral | 0.008829 | 0.03849 | 539.1532 | 0.229395 | 0.818649 | N/A |
| Condition:Aversive | 0.018929 | 0.038413 | 539.5451 | 0.492784 | 0.622365 | 0.918928 |
| TI:Neutral | 0.023194 | 0.038687 | 539.2538 | 0.599535 | 0.549068 | N/A |
| TI:Aversive | 0.0187 | 0.037655 | 539.352 | 0.496601 | 0.619673 | 0.918928 |
| Condition:Unguided dmPFC Site | 0.004823 | 0.04815 | 539.5003 | 0.10017 | 0.920247 | N/A |
| TI:Unguided dmPFC Site | -0.03134 | 0.044666 | 539.2279 | -0.7016 | 0.483229 | N/A |
| **Neutral:Unguided dmPFC Site** | **0.093186** | **0.044972** | **539.0657** | **2.072095** | **0.038731*** | N/A |
| Aversive:Unguided dmPFC Site | -0.00842 | 0.047981 | 539.4871 | -0.17558 | 0.860693 | 0.918928 |
| Condition:TI:Neutral | 0.004897 | 0.054331 | 539.5467 | 0.090141 | 0.928209 | N/A |
| Condition:TI:Aversive | -0.00949 | 0.054385 | 539.6729 | -0.17455 | 0.861495 | 0.918928 |
| Condition:TI:Unguided dmPFC Site | 0.035042 | 0.066604 | 539.5395 | 0.526125 | 0.599018 | N/A |
| Condition:Neutral:Unguided dmPFC Site | -0.01619 | 0.066893 | 539.2345 | -0.24198 | 0.808886 | N/A |
| Condition:Aversive:Unguided dmPFC Site | 0.046052 | 0.068099 | 539.8895 | 0.676253 | 0.499169 | 0.918928 |
| TI:Neutral:Unguided dmPFC Site | -0.09251 | 0.064455 | 539.1276 | -1.43526 | 0.151794 | N/A |
| TI:Aversive:Unguided dmPFC Site | 0.077728 | 0.067379 | 539.4915 | 1.153582 | 0.249183 | 0.724896 |
| Condition:TI:Neutral:Unguided dmPFC Site | -0.05112 | 0.094066 | 539.3343 | -0.54344 | 0.587055 | N/A |
| Condition:TI:Aversive:Unguided dmPFC Site | -0.14738 | 0.095315 | 539.8035 | -1.54621 | 0.122639 | 0.490556 |

**Color-Word Stroop: Stimulation Time**

(Omnibus: full factorial model testing Color-Word Stroop response time based on stimulation time)

| RT~Condition*Stimulation Time*Stimulation Site+Run Order+(1\|Subject) | | |  |  |  |  |
| --- | --- | --- | --- | --- | --- | --- |
|  | Estimate | Error | df | t | Pr(>\|t\|) | FDR |
| **(Intercept)** | **0.64964** | **0.0462** | **28.40178** | **14.06157** | **2.53E-14***** | **2.87E-13** |
| **Condition** | **0.059983** | **0.010251** | **9432.865** | **5.851272** | **5.04E-09***** | **3.43E-08** |
| **Stimulation Time** | **0.003508** | **0.001146** | **7284.365** | **3.06131** | **0.002212**** | **0.009401** |
| **rmPFC Site** | **0.028754** | **0.011029** | **9439.193** | **2.607171** | **0.009144**** | **0.03109** |
| **Unguided dmPFC Site** | **0.165683** | **0.057749** | **36.34866** | **2.869039** | **0.006818**** | **0.025757** |
| Run Order 2 | -0.00712 | 0.060241 | 27.05859 | -0.1182 | 0.906786 | 0.934264 |
| Run Order 3 | 0.044799 | 0.064205 | 31.11198 | 0.69775 | 0.490516 | 0.667102 |
| **Condition:Stimulation Time** | **0.001368** | **0.000588** | **9432.87** | **2.325286** | **0.020078**** | **0.056888** |
| Condition:rmPFC Site | 0.009088 | 0.015321 | 9432.888 | 0.593185 | 0.553071 | 0.723247 |
| Condition:Unguided dmPFC Site | 0.015452 | 0.016102 | 9432.862 | 0.959645 | 0.337258 | 0.546037 |
| Stimulation Time:rmPFC Site | -0.00098 | 0.000607 | 9436.053 | -1.61195 | 0.107006 | 0.214012 |
| **Stimulation Time:Unguided dmPFC Site** | **-0.0102** | **0.001667** | **3444.057** | **-6.11682** | **1.06E-09***** | **9.01E-09** |
| Condition:Stimulation Time:rmPFC Site | -0.00126 | 0.000854 | 9432.897 | -1.4744 | 0.140406 | 0.265211 |
| **Condition:Stimulation Time:Unguided dmPFC Site** | **-0.00293** | **0.000927** | **9432.861** | **-3.1655** | **0.001553***** | **0.007543** |

**Color-Word Stroop: Stimulation Time**

(Omnibus; Relevel: Unguided dmPFC)

| RT~Condition*Stimulation Time*Stimulation Site+Run Order+(1\|Subject) | | |  |  |  |  |
| --- | --- | --- | --- | --- | --- | --- |
|  | Estimate | Error | df | t | Pr(>\|t\|) | FDR |
| **(Intercept)** | **0.815324** | **0.055057** | **30.23403** | **14.80863** | **2.13E-15***** | N/A |
| **Condition** | **0.075435** | **0.012417** | **9432.858** | **6.075312** | **1.29E-09***** | N/A |
| **Stimulation Time** | **-0.00669** | **0.001251** | **5874.138** | **-5.3483** | **9.21E-08***** | N/A |
| **dmPFC Site** | **-0.16568** | **0.057749** | **36.3487** | **-2.86904** | **0.006818**** | N/A |
| **rmPFC Site** | **-0.13693** | **0.057906** | **36.74137** | **-2.36468** | **0.023442*** | **0.06131** |
| Run Order 2 | -0.00712 | 0.060241 | 27.0586 | -0.1182 | 0.906786 | N/A |
| Run Order 3 | 0.044799 | 0.064205 | 31.112 | 0.69775 | 0.490516 | N/A |
| **Condition:Stimulation Time** | **-0.00157** | **0.000716** | **9432.858** | **-2.18656** | **0.028799*** | N/A |
| Condition:dmPFC Site | -0.01545 | 0.016102 | 9432.862 | -0.95965 | 0.337258 | N/A |
| Condition:rmPFC Site | -0.00636 | 0.016848 | 9432.902 | -0.37772 | 0.705649 | 0.783479 |
| **Stimulation Time:dmPFC Site** | **0.010197** | **0.001667** | **3444.057** | **6.116818** | **1.06E-09***** | N/A |
| **Stimulation Time:rmPFC Site** | **0.009219** | **0.001679** | **3480.995** | **5.489869** | **4.31E-08***** | **2.44E-07** |
| **Condition:Stimulation Time:dmPFC Site** | **0.002934** | **0.000927** | **9432.861** | **3.1655** | **0.001553**** | N/A |
| **Condition:Stimulation Time:rmPFC Site** | **0.001675** | **0.000946** | **9432.881** | **1.769984** | **0.076762*** | **0.163119** |

**Color-Word Stroop: Stimulation Time – Congruent Trials**

(Deconstruction of Stimulation Time)

| RT~Stimulation Time*Stimulation Site+Run Order+(1\|Subject) | | | |  |  |
| --- | --- | --- | --- | --- | --- |
|  | Estimate | Std.Error | df | t | Pr(>\|t\|) |
| **(Intercept)** | **0.639246** | **0.05059** | **36.17752** | **12.63584** | **7.83E-15***** |
| **Stimulation Time** | **0.003814** | **0.001492** | **2870.67** | **2.555647** | **0.01065**** |
| **Unguided dmPFC Site** | **0.153609** | **0.056716** | **43.04427** | **2.708394** | **0.009662**** |
| **rmPFC Site** | **0.034006** | **0.0111** | **4781.277** | **3.063735** | **0.002198**** |
| Run Order 1 | 0.010761 | 0.056199 | 28.47306 | 0.191475 | 0.849512 |
| Run Order 3 | 0.043575 | 0.060186 | 28.2356 | 0.723998 | 0.475023 |
| **Stimulation Time:Unguided dmPFC Site** | **-0.00939** | **0.002115** | **873.703** | **-4.43816** | **1.02E-05***** |
| **Stimulation Time:rmPFC Site** | **-0.00117** | **0.000603** | **4775.427** | **-1.93769** | **0.05272^.^** |

**Color-Word Stroop: Stimulation Time – Incongruent Trials**

(Deconstruction: Stimulation Time)

| RT~Stimulation Time*Stimulation Site+Run Order+(1\|Subject) | | | |  |  |
| --- | --- | --- | --- | --- | --- |
|  | Estimate | Std.Error | df | t | Pr(>\|t\|) |
| **(Intercept)** | **0.710634** | **0.059213** | **34.07373** | **12.00136** | **8.69E-14***** |
| **Stimulation Time** | **0.004139** | **0.001584** | **3270.965** | **2.613017** | **0.009016**** |
| **Unguided dmPFC Site** | **0.176856** | **0.06556** | **40.03025** | **2.69764** | **0.010172*** |
| **rmPFC Site** | **0.033464** | **0.011617** | **4645.833** | **2.880679** | **0.003986**** |
| Run Order 1 | 0.004409 | 0.066566 | 27.93492 | 0.066239 | 0.947659 |
| Run Order 3 | 0.063728 | 0.071357 | 27.83272 | 0.893084 | 0.379469 |
| **Stimulation Time:Unguided dmPFC Site** | **-0.01278** | **0.002236** | **1296.635** | **-5.71576** | **1.35E-08***** |
| **Stimulation Time:rmPFC Site** | **-0.00211** | **0.000628** | **4640.968** | **-3.35299** | **0.000806***** |

**Color-Word Stroop: Stimulation Time – Congruent Trials**

(Deconstruction: Stimulation Time; Relevel: Unguided dmPFC)

| RT~Stimulation Time*Stimulation Site+Run Order+(1\|Subject) | | | | | |
| --- | --- | --- | --- | --- | --- |
|  | Estimate | Std.Error | df | t | Pr(>\|t\|) |
| **(Intercept)** | **0.792855** | **0.052859** | **41.67495** | **14.99952** | **2.13E-18***** |
| **Stimulation Time** | **-0.00557** | **0.001621** | **1878.882** | **-3.4369** | **0.000601***** |
| **dmPFC Site** | **-0.15361** | **0.056716** | **43.04424** | **-2.70839** | **0.009662**** |
| rmPFC Site | -0.1196 | 0.056898 | 43.58191 | -2.10206 | 0.041362 |
| Run Order 1 | 0.010761 | 0.056199 | 28.47304 | 0.191475 | 0.849512 |
| Run Order 3 | 0.043575 | 0.060186 | 28.23558 | 0.723998 | 0.475023 |
| **Stimulation Time:dmPFC Site** | **0.009386** | **0.002115** | **873.703** | **4.438156** | **1.02E-05***** |
| **Stimulation Time:rmPFC Site** | **0.008218** | **0.002127** | **882.6062** | **3.863679** | **0.00012***** |

**Color-Word Stroop: Stimulation Time – Incongruent Trials**

(Deconstruction: Stimulation Time; Relevel: Unguided dmPFC relevel)

| RT~Stimulation Time*Stimulation Site+Run Order+(1\|Subject) | | | | | |
| --- | --- | --- | --- | --- | --- |
|  | Estimate | Std.Error | df | t | Pr(>\|t\|) |
| **(Intercept)** | **0.710634** | **0.059213** | **34.07373** | **12.00136** | **8.69E-14***** |
| **Stimulation Time** | **0.004139** | **0.001584** | **3270.965** | **2.613017** | **0.009016**** |
| **Unguided dmPFC Site** | **0.176856** | **0.06556** | **40.03025** | **2.69764** | **0.010172**** |
| **rmPFC Site** | **0.033464** | **0.011617** | **4645.833** | **2.880679** | **0.003986**** |
| Run Order 1 | 0.004409 | 0.066566 | 27.93492 | 0.066239 | 0.947659 |
| Run Order 3 | 0.063728 | 0.071357 | 27.83272 | 0.893084 | 0.379469 |
| **Stimulation Time:Unguided dmPFC Site** | **-0.01278** | **0.002236** | **1296.635** | **-5.71576** | **1.35E-08***** |
| **Stimulation Time:rmPFC Site** | **-0.00211** | **0.000628** | **4640.968** | **-3.35299** | **0.000806***** |

**Color-Word Stroop: Stimulation Time – rmPFC Site**

(Deconstruction: Stimulation Time)

| RT~Condition*Stimulation Time+Run Order+(1\|Subject) | |  |  |  |  |
| --- | --- | --- | --- | --- | --- |
|  | Estimate | Error | df | t | Pr(>\|t\|) |
| **(Intercept)** | **0.744017** | **0.055694** | **14.6291** | **13.35907** | **1.35E-09***** |
| **Condition** | **0.069397** | **0.011743** | **3088.077** | **5.909854** | **3.80E-09***** |
| **Stimulation Time** | **0.00473** | **0.001793** | **3076.968** | **2.638548** | **0.008368**** |
| Run Order 2 | -0.14484 | 0.089653 | 15.70029 | -1.61561 | 0.126092 |
| Run Order 3 | -0.08835 | 0.084839 | 22.69984 | -1.04135 | 0.308678 |
| Condition:Stimulation Time | 0.000102 | 0.000638 | 3088.053 | 0.159647 | 0.873169 |

**Color-Word Stroop: Stimulation Time – dmPFC Site**

(Deconstruction: Stimulation Time)

| RT~Condition*Stimulation Time+Run Order+(1\|Subject) | |  |  |  |  |
| --- | --- | --- | --- | --- | --- |
|  | Estimate | Error | df | t | Pr(>\|t\|) |
| **(Intercept)** | **0.695771** | **0.049431** | **16.63657** | **14.07553** | **1.15E-10***** |
| **Condition** | **0.059563** | **0.010105** | **3602.028** | **5.894543** | **4.10E-09***** |
| **Stimulation Time** | **0.002501** | **0.001462** | **3600.996** | **1.710722** | **0.087219^.^** |
| Run Order 2 | -0.07996 | 0.077477 | 17.41953 | -1.032 | 0.316188 |
| Run Order 3 | 0.010524 | 0.076242 | 23.44427 | 0.138035 | 0.891392 |
| **Condition:Stimulation Time** | **0.001384** | **0.00058** | **3602.034** | **2.387517** | **0.017014^*^** |

**Color-Word Stroop: Stimulation Time – Unguided dmPFC Site**

(Deconstruction: Stimulation Time)

| RT~Condition*Stimulation Time+Run Order+(1\|Subject) | |  |  |  |  |
| --- | --- | --- | --- | --- | --- |
|  | Estimate | Error | df | t | Pr(>\|t\|) |
| **(Intercept)** | **0.742127** | **0.067427** | **8.229457** | **11.00635** | **3.31E-06***** |
| **Condition** | **0.075425** | **0.011973** | **2727.012** | **6.299802** | **3.46E-10***** |
| **Stimulation Time** | **-0.00708** | **0.001231** | **2728.872** | **-5.75242** | **9.78E-09***** |
| Run Order 2 | 0.115186 | 0.091021 | 8.432291 | 1.26549 | 0.239542 |
| Run Order 3 | 0.172558 | 0.119272 | 8.946991 | 1.446754 | 0.182075 |
| **Condition:Stimulation Time** | **-0.00157** | **0.000691** | **2727.013** | **-2.26765** | **0.023428*** |

**Color-Word Stroop: Stimulation Time – dmPFC Site, Congruent Trials**

(Deconstruction: Stimulation Time)

| RT~Stimulation Time+Run Order+(1\|Subject) |  |  |  |  |  |
| --- | --- | --- | --- | --- | --- |
|  | Estimate | Error | df | t | Pr(>\|t\|) |
| **(Intercept)** | **0.688117** | **0.0431** | **16.72179** | **15.96575** | **1.49E-11***** |
| **Stimulation Time** | **0.003628** | **0.001901** | **1812.051** | **1.908698** | **0.056459^.^** |
| Run Order 2 | -0.085 | 0.069316 | 19.3514 | -1.22633 | 0.234789 |
| Run Order 3 | -0.01126 | 0.073657 | 34.88387 | -0.1529 | 0.87936 |

**Color-Word Stroop: Stimulation Time – dmPFC Site, Incongruent Trials**

(Deconstruction: Stimulation Time)

| RT~Stimulation Time+Run Order+(1\|Subject) |  |  |  |  |  |
| --- | --- | --- | --- | --- | --- |
|  | Estimate | Error | df | t | Pr(>\|t\|) |
| **(Intercept)** | **0.761768** | **0.057455** | **16.49328** | **13.25855** | **3.22E-10***** |
| Stimulation Time | 0.003134 | 0.002109 | 1762.726 | 1.486084 | 0.137436 |
| Run Order 2 | -0.07963 | 0.091337 | 18.25642 | -0.87182 | 0.394629 |
| Run Order 3 | 0.024884 | 0.093296 | 28.36 | 0.266725 | 0.791611 |

**Color-Word Stroop: Stimulation Time – rmPFC Site, Congruent Trials**

(Deconstruction: Stimulation Time)

| RT~Stimulation Time+Run Order+(1\|Subject) |  |  |  |  |  |
| --- | --- | --- | --- | --- | --- |
|  | Estimate | Error | df | t | Pr(>\|t\|) |
| **(Intercept)** | **0.74138** | **0.051082** | **14.69764** | **14.51363** | **4.08E-10***** |
| **Stimulation Time** | **0.004488** | **0.00243** | **1529.291** | **1.846876** | **0.064958^.^** |
| Run Order 2 | -0.12982 | 0.084942 | 17.88207 | -1.52837 | 0.143916 |
| Run Order 3 | -0.08138 | 0.087689 | 35.81837 | -0.92807 | 0.359584 |

**Color-Word Stroop: Stimulation Time – rmPFC Site, Incongruent Trials**

(Deconstruction: Stimulation Time)

| RT~Stimulation Time+Run Order+(1\|Subject) | Estimate | Error | df | t | Pr(>\|t\|) |
| --- | --- | --- | --- | --- | --- |
| **(Intercept)** | **0.816273** | **0.061558** | **14.4892** | **13.26022** | **1.68E-09***** |
| **Stimulation Time** | **0.005042** | **0.002501** | **1514.846** | **2.015925** | **0.043984*** |
| Run Order 2 | -0.16001 | 0.101156 | 16.85398 | -1.58183 | 0.132271 |
| Run Order 3 | -0.09466 | 0.100422 | 29.23106 | -0.94259 | 0.353614 |

**Color-Word Stroop: Stimulation Time – Unguided dmPFC, Congruent Trials**

(Deconstruction: Stimulation Time)

| RT~Stimulation Time+Run Order+(1\|Subject) |  |  |  |  |  |
| --- | --- | --- | --- | --- | --- |
|  | Estimate | Error | df | t | Pr(>\|t\|) |
| **(Intercept)** | **0.742231** | **0.06679** | **8.205959** | **11.11283** | **3.14E-06***** |
| **Stimulation Time** | **-0.00634** | **0.001676** | **1363.412** | **-3.78282** | **0.000162***** |
| Run Order 2 | 0.101279 | 0.091454 | 8.893702 | 1.107429 | 0.297159 |
| Run Order 3 | 0.151448 | 0.121572 | 9.981777 | 1.245741 | 0.241306 |

**Color-Word Stroop: Stimulation Time – Unguided dmPFC Site, Incongruent Trials**

(Deconstruction: Stimulation Time)

| RT~Stimulation Time+Run Order+(1\|Subject) |  |  |  |  |  |
| --- | --- | --- | --- | --- | --- |
|  | Estimate | Error | df | t | Pr(>\|t\|) |
| **(Intercept)** | **0.817041** | **0.06914** | **8.189488** | **11.81713** | **1.98E-06***** |
| **Stimulation Time** | **-0.00929** | **0.00165** | **1353.905** | **-5.63046** | **2.18E-08***** |
| Run Order 2 | 0.12794 | 0.094466 | 8.800871 | 1.354343 | 0.209374 |
| Run Order 3 | 0.191673 | 0.125251 | 9.778617 | 1.530306 | 0.157625 |

**Color-Word Stroop: Accuracy**

(Omnibus: full factorial model testing Color-Word Stroop accuracy based on stimulation time)

| Correct~Condition*Stimulation Time*Stimulation Site+Run Order+(1\|Subject) | | |  |  |  |
| --- | --- | --- | --- | --- | --- |
|  | Estimate | Error | z | Pr(>\|z\|) | FDR |
| **(Intercept)** | **4.134331** | **0.374658** | **11.03494** | **2.59E-28***** | **4.66E-27** |
| **Condition** | **-0.71212** | **0.309349** | **-2.30199** | **0.021336*** | **0.123341** |
| **Stimulation Time** | **-0.08449** | **0.029231** | **-2.89046** | **0.003847**** | **0.034623** |
| Unguided dmPFC Site | -0.72063 | 0.573884 | -1.25571 | 0.209221 | 0.376598 |
| rmPFC Site | 0.244547 | 0.36564 | 0.668818 | 0.503611 | 0.566562 |
| **Run Order 2** | **0.839978** | **0.4459** | **1.883781** | **0.059595^.^** | **0.153244** |
| **Run Order 3** | **1.29975** | **0.655436** | **1.98303** | **0.047364*** | **0.142092** |
| **Condition:Stimulation Time** | **0.036907** | **0.017467** | **2.11295** | **0.034605*** | **0.124578** |
| Condition:Unguided dmPFC Site | 0.19725 | 0.460029 | 0.428777 | 0.668085 | 0.668085 |
| Condition:rmPFC Site | -0.47144 | 0.458777 | -1.02761 | 0.304134 | 0.437993 |
| **Stimulation Time:Unguided dmPFC Site** | **0.074149** | **0.033618** | **2.205642** | **0.027409*** | **0.123341** |
| Stimulation Time:rmPFC Site | -0.01318 | 0.018534 | -0.71095 | 0.477114 | 0.566562 |
| Condition:Stimulation Time:Unguided dmPFC Site | -0.0291 | 0.029921 | -0.97252 | 0.33079 | 0.437993 |
| Condition:Stimulation Time:rmPFC Site | 0.024051 | 0.02524 | 0.952859 | 0.340661 | 0.437993 |

**Color-Word Stroop: Accuracy**

(Omnibus; Relevel: Unguided dmPFC)

| Correct~Congruent*StimTime3*CongSite+RunOrder+(1\|Sub) | |  |  |  |  |
| --- | --- | --- | --- | --- | --- |
|  | Estimate | Std.Error | z | Pr(>\|z\|) | FDR |
| **(Intercept)** | **2.898533** | **0.426226** | **6.800452** | **1.04e-11** | N/A |
| Condition | 0.515782 | 0.34024 | 1.515936 | 0.129536 | N/A |
| Stimulation Time | -0.0025 | 0.029934 | -0.08357 | 0.933402 | N/A |
| dmPFC Site | 0.523287 | 0.536798 | 0.974831 | 0.329644 | N/A |
| rmPFC Site | 0.296951 | 0.539624 | 0.550293 | 0.582118 | .61636 |
| **Run Order 2** | **0.839962** | **0.445997** | **1.883336** | **0.059655** | N/A |
| **Run Order 3** | **1.299842** | **0.655647** | **1.982533** | **0.04742** | N/A |
| Condition:Stimulation Tme | -0.00787 | 0.024284 | -0.3239 | 0.746017 | N/A |
| Condition:dmPFC Site | 0.197299 | 0.459611 | 0.429273 | 0.667725 | N/A |
| Condition:rmPFC Site | 0.666644 | 0.48076 | 1.386645 | 0.16555 | .3311 |
| Stimulation Time:dmPFC Site | -0.04507 | 0.031968 | -1.40975 | 0.158614 | N/A |
| Stimulation Time:rmPFC Site | -0.03422 | 0.032088 | -1.06644 | 0.286224 | .437993 |
| Condition:Stimulation Time:dmPFC Site | -0.02909 | 0.029903 | -0.97284 | 0.330631 | N/A |
| **Condition:Stimulation Time:rmPFC Site** | **-0.05303** | **0.0304** | **-1.7445** | **0.081072** | **.182411** |

**Color-Word: Accuracy – dmPFC Site**

(Deconstruction: Accuracy)

| Correct~Condition*Stimulation Time+Run Order+(1\|Subject) | |  |  |  |
| --- | --- | --- | --- | --- |
|  | Estimate | Error | z | Pr(>\|z\|) |
| **(Intercept)** | **4.149118** | **0.399854** | **10.37658** | **3.17E-25***** |
| **Condition** | **-0.71205** | **0.308045** | **-2.31151** | **0.020804*** |
| **Stimulation Time** | **-0.10033** | **0.042149** | **-2.38027** | **0.0173*** |
| Run Order 2 | 0.997701 | 0.678667 | 1.470089 | 0.141538 |
| Run Order 3 | 1.719378 | 1.019901 | 1.685829 | 0.091829 |
| **Condition:Stimulation Time** | **0.037029** | **0.017491** | **2.116986** | **0.034261*** |

**Color-Word: Accuracy – rmPFC Site**

(Deconstruction: Accuracy)

| Correct~Condition*Stimulation Time+Run Order+(1\|Subject) | |  |  |  |
| --- | --- | --- | --- | --- |
|  | Estimate | Error | z | Pr(>\|z\|) |
| **(Intercept)** | **4.333719** | **0.46391** | **9.341731** | **9.48E-21***** |
| **Condition** | **-1.16917** | **0.338006** | **-3.45901** | **0.000542***** |
| **Stimulation Time** | **-0.0771** | **0.046833** | **-1.64636** | **0.099691^.^** |
| Run Order 2 | 0.46538 | 0.810248 | 0.574368 | 0.565719 |
| Run Order 3 | 0.813824 | 1.148104 | 0.708842 | 0.478422 |
| **Condition:Stimulation Time** | **0.059981** | **0.018144** | **3.30587** | **0.000947***** |

**Color-Word: Accuracy – Unguided dmPFC Site**

(Deconstruction: Accuracy)

| Correct~Condition*Stimulation Time+Run Order+(1\|Subject) | |  |  |  |
| --- | --- | --- | --- | --- |
|  | Estimate | Error | z | Pr(>\|z\|) |
| **(Intercept)** | **3.379572** | **0.44045** | **7.673007** | **1.68E-14***** |
| Condition | -0.51574 | 0.341361 | -1.51082 | 0.130834 |
| Stimulation Time | -0.01612 | 0.038867 | -0.41481 | 0.678281 |
| Run Order 2 | 0.966546 | 0.656467 | 1.472346 | 0.140927 |
| Run Order 3 | 1.532029 | 1.110894 | 1.379095 | 0.167865 |
| Condition:Stimulation Time | 0.00792 | 0.024413 | 0.324404 | 0.745632 |

**Color-Word: Accuracy – dmPFC Site, Congruent Trials**

(Deconstruction: Accuracy)

| Correct~Stimulation Time+Run Order+(1\|Subject) |  |  |  |  |
| --- | --- | --- | --- | --- |
|  | Estimate | Error | z | Pr(>\|z\|) |
| **(Intercept)** | **3.838419** | **0.374295** | **10.25505** | **1.12E-24***** |
| Stimulation Time | -0.07598 | 0.053814 | -1.41196 | 0.157963 |
| Run Order 2 | 0.875883 | 0.801764 | 1.092445 | 0.274637 |
| Run Order 3 | 1.348942 | 1.284424 | 1.050231 | 0.293612 |

**Color-Word: Accuracy – dmPFC Site, Incongruent Trials**

(Deconstruction: Accuracy)

| Correct~Stimulation Time+Run Order+(1\|Subject) |  |  |  |  |
| --- | --- | --- | --- | --- |
|  | Estimate | Error | z | Pr(>\|z\|) |
| **(Intercept)** | **3.673332** | **0.449607** | **8.170093** | **3.08E-16***** |
| **Stimulation Time** | **-0.10913** | **0.055661** | **-1.96061** | **0.049925*** |
| **Run Order 2** | **1.48864** | **0.873546** | **1.704135** | **0.088356^.^** |
| **Run Order 3** | **2.870501** | **1.35857** | **2.112884** | **0.034611*** |

**Color-Word: Accuracy – rmPFC site, Congruent Trials**

(Deconstruction: Accuracy)

| Correct~Stimulation Time+Run Order+(1\|Subject) |  |  |  |  |
| --- | --- | --- | --- | --- |
|  | Estimate | Error | z | Pr(>\|z\|) |
| **(Intercept)** | **3.988905** | **0.387694** | **10.28879** | **7.92E-25***** |
| **Stimulation Time** | **-0.14975** | **0.061109** | **-2.45056** | **0.014263**** |
| **Run Order 2** | **2.161675** | **0.962354** | **2.246236** | **0.024689*** |
| **Run Order 3** | **2.984787** | **1.430338** | **2.08677** | **0.036909*** |

**Color-Word: Accuracy – rmPFC site, Incongruent Trials**

(Deconstruction: Accuracy)

| Correct~Stimulation Time+Run Order+(1\|Subject) |  |  |  |  |
| --- | --- | --- | --- | --- |
|  | Estimate | Error | z | Pr(>\|z\|) |
| **(Intercept)** | **3.378796** | **0.55004** | **6.142819** | **8.11E-10** |
| Stimulation Time | 0.026175 | 0.062349 | 0.419821 | 0.674616 |
| Run Order 2 | -0.47376 | 1.098945 | -0.43111 | 0.666392 |
| Run Order 3 | -0.28515 | 1.565658 | -0.18213 | 0.85548 |

**Color-Word: Accuracy – unguided dmPFC site, Congruent Trials**

(Deconstruction: Accuracy)

| Correct~Stimulation Time+Run Order+(1\|Subject) |  |  |  |  |
| --- | --- | --- | --- | --- |
|  | Estimate | Error | z | Pr(>\|z\|) |
| **(Intercept)** | **3.264422** | **0.390304** | **8.363804** | **6.07E-17***** |
| Stimulation Time | -0.01134 | 0.050466 | -0.22464 | 0.822257 |
| Run Order 2 | 0.698278 | 0.75051 | 0.930405 | 0.352161 |
| Run Order 3 | 1.959294 | 1.441254 | 1.359437 | 0.174008 |

**Color-Word: Accuracy – unguided dmPFC site, Incongruent Trials**

(Deconstruction: Accuracy)

| Correct~Stimulation Time+Run Order+(1\|Subject) |  |  |  |  |
| --- | --- | --- | --- | --- |
|  | Estimate | Error | z | Pr(>\|z\|) |
| **(Intercept)** | **2.764239** | **0.449967** | **6.143205** | **8.09E-10***** |
| Stimulation Time | 0.014904 | 0.045759 | 0.325713 | 0.744641 |
| Run Order 2 | 0.927649 | 0.792608 | 1.170375 | 0.24185 |
| Run Order 3 | 0.648669 | 1.327091 | 0.488791 | 0.62499 |

**Affective Stroop: Stimulation Time**

(Omnibus: full factorial model testing Affective Stroop response time based on stimulation time)

| RT~Condition*Stimulation Time*Valence+Stimulation Site+Run Order+(1\|Subject) | | | | |  |  |
| --- | --- | --- | --- | --- | --- | --- |
|  | Estimate | Error | df | t | Pr(>\|t\|) | FDR |
| **(Intercept)** | **0.590711** | **0.027995** | **29.15908** | **21.10091199** | **3.29E-19***** | **4.19E-18** |
| **Condition** | **0.075181** | **0.009171** | **5647.245** | **8.19774211** | **3.00E-16***** | **3.06E-15** |
| **Stimulation Time** | **-0.00774** | **0.000701** | **5672.731** | **-11.04378086** | **4.53E-28***** | **1.16E-26** |
| Aversive | 0.005399 | 0.009307 | 5647.192 | 0.580086911 | 0.561879 | 0.91158 |
| Pleasant | -0.01125 | 0.00923 | 5647.174 | -1.219198755 | 0.22282 | 0.568191 |
| Unguided dmPFC Site | 0.015528 | 0.029445 | 26.00337 | 0.527340787 | 0.602428 | 0.91158 |
| **Run Order 2** | **0.119399** | **0.035108** | **28.66595** | **3.400959136** | **0.001996**** | **0.014542** |
| **Run Order 3** | **0.214553** | **0.039006** | **38.58978** | **5.500530662** | **2.64E-06***** | **2.24E-05** |
| Condition:Stimulation Time | 0.000603 | 0.000455 | 5647.371 | 1.324330519 | 0.185447 | 0.525433 |
| Condition:Aversive | 0.015438 | 0.013009 | 5647.461 | 1.186722378 | 0.235387 | 0.571654 |
| **Condition:Pleasant** | **0.024242** | **0.012997** | **5647.35** | **1.865171927** | **0.062209*** | **0.279008** |
| Stimulation Time:Aversive | 0.000376 | 0.00046 | 5647.189 | 0.81632394 | 0.414349 | 0.812762 |
| **Stimulation Time:Pleasant** | **0.001277** | **0.000454** | **5647.239** | **2.811826813** | **0.004943**** | **0.02801** |
| **Condition:Stimulation Time:Aversive** | **-0.00108** | **0.000646** | **5647.501** | **-1.666236768** | **0.095722^.^** | **0.348702** |
| **Condition:Stimulation Time:Pleasant** | **-0.00188** | **0.000644** | **5647.524** | **-2.928052344** | **0.003425**** | **0.021834** |

**Affective Stroop: Stimulation Time**

(Relevel: Aversive Trials)

| RT~Condition*Stimulation Time*Valence+Stimulation Site+Run Order+(1\|Subject) | | | | |  |  |
| --- | --- | --- | --- | --- | --- | --- |
|  | Estimate | Error | df | t | Pr(>\|t\|) | FDR |
| **(Intercept)** | **0.59611** | **0.028** | **29.18241** | **21.28939269** | **2.52E-19***** | N/A |
| **Condition** | **0.090619** | **0.009216** | **5647.331** | **9.833319964** | **1.23E-22***** | N/A |
| **Stimulation Time** | **-0.00737** | **0.000706** | **5672.71** | **-10.43347684** | **2.95E-25***** | N/A |
| Neutral | -0.0054 | 0.009307 | 5647.192 | -0.580086902 | 0.561879 | N/A |
| **Pleasant** | **-0.01665** | **0.009236** | **5647.169** | **-1.802967115** | **0.071447^.^** | 0.280292 |
| Unguided dmPFC Site | 0.015528 | 0.029445 | 26.00341 | 0.527341055 | 0.602428 | N/A |
| **Run Order 2** | **0.119399** | **0.035108** | **28.66599** | **3.400960695** | **0.001996**** | N/A |
| **Run Order 3** | **0.214553** | **0.039006** | **38.58985** | **5.500532812** | **2.64E-06***** | N/A |
| Condition:Stimulation Time | -0.00047 | 0.000457 | 5647.364 | -1.03401164 | 0.301175 | N/A |
| Condition:Neutral | -0.01544 | 0.013009 | 5647.461 | -1.186722386 | 0.235387 | N/A |
| Condition:Pleasant | 0.008804 | 0.013034 | 5647.469 | 0.675483733 | 0.499396 | 0.878248 |
| Stimulation Time:Neutral | -0.00038 | 0.00046 | 5647.189 | -0.816323946 | 0.414349 | N/A |
| **Stimulation Time:Pleasant** | **0.000901** | **0.000461** | **5647.282** | **1.953487666** | **0.050811*** | 0.259136 |
| **Condition:Stimulation Time:Neutral** | **0.001076** | **0.000646** | **5647.501** | **1.666236778** | **0.095722^.^** | N/A |
| Condition:Stimulation Time:Pleasant | -0.00081 | 0.000645 | 5647.542 | -1.253285235 | 0.210154 | 0.564098 |

**Affective Stroop: Stimulation Time - Emotional Stimuli**

(Alternative Omnibus: full factorial model testing Affective Stroop response time based on stimulation time; emotional stimuli pooled)

| RT~Condition*Stimulation Time*Valence2+Stimulation Site+Run Order+(1\|Subject) | | | | |  |
| --- | --- | --- | --- | --- | --- |
|  | Estimate | Std.Error | df | t | Pr(>\|t\|) |
| **(Intercept)** | **0.587655** | **0.027604** | **27.57651** | **21.28894** | **1.18E-18***** |
| **Condition** | **0.095226** | **0.006504** | **5651.108** | **14.64188** | **1.12E-47***** |
| **Stimulation Time** | **-0.0069** | **0.000665** | **5675.447** | **-10.3783** | **5.21E-25***** |
| Neutral | 0.003114 | 0.008036 | 5651.187 | 0.387471 | 0.698422 |
| Unguided dmPFC Site | 0.015375 | 0.029442 | 26.00259 | 0.522218 | 0.605939 |
| **Run Order 2** | **0.119339** | **0.035104** | **28.66663** | **3.399593** | **0.002003**** |
| **Run Order 3** | **0.214494** | **0.039003** | **38.59487** | **5.499434** | **2.64E-06***** |
| **Condition:Stimulation Time** | **-0.00089** | **0.000322** | **5651.112** | **-2.7721** | **0.005588**** |
| **Condition:Neutral** | **-0.02004** | **0.011253** | **5651.385** | **-1.78066** | **0.075022^.^** |
| **Stimulation Time:Neutral** | **-0.00084** | **0.000395** | **5651.192** | **-2.12328** | **0.033774*** |
| **Condition:Stimulation Time:Neutral** | **0.001494** | **0.000558** | **5651.506** | **2.677159** | **0.007446**** |

**Affective Stroop: Stimulation Time – Neutral Stimuli**

(Deconstruction: Stimulation Time)

| RT~Condition*Stimulation Time+Stimulation Site+Run Order+(1\|Subject) | | | | |  |
| --- | --- | --- | --- | --- | --- |
|  | Estimate | Error | df | t | Pr(>\|t\|) |
| **(Intercept)** | **0.587315** | **0.026535** | **29.3668** | **22.13318321** | **7.22E-20***** |
| **Condition** | **0.074657** | **0.008962** | **1866.729** | **8.330340013** | **1.54E-16***** |
| **Stimulation Time** | **-0.00634** | **0.001086** | **1889.02** | **-5.838685983** | **6.18E-09***** |
| Unguided dmPFC Site | 0.020649 | 0.027847 | 25.97824 | 0.741530173 | 0.465023 |
| **Run Order 2** | **0.094287** | **0.03482** | **34.44929** | **2.707807672** | **0.010469*** |
| **Run Order 3** | **0.17417** | **0.043344** | **71.35403** | **4.01833289** | **0.000143***** |
| Condition:Stimulation Time | 0.000647 | 0.000445 | 1867.126 | 1.4517291 | 0.146745 |

**Affective Stroop: Stimulation Time – Pleasant Stimuli**

(Deconstruction: Stimulation Time)

| RT~Condition*Stimulation Time+Stimulation Site+Run Order+(1\|Subject) | | | | |  |
| --- | --- | --- | --- | --- | --- |
|  | Estimate | Error | df | t | Pr(>\|t\|) |
| **(Intercept)** | **0.583289** | **0.029486** | **28.97342** | **19.78215645** | **2.26E-18***** |
| **Condition** | **0.098424** | **0.009305** | **1875.721** | **10.57703625** | **1.95E-25***** |
| **Stimulation Time** | **-0.00722** | **0.001128** | **1900.392** | **-6.403540748** | **1.91E-10***** |
| Unguided dmPFC Site | 0.00784 | 0.031073 | 26.05924 | 0.252305847 | 0.802784 |
| **Run Order 2** | **0.133474** | **0.038532** | **33.4748** | **3.463951144** | **0.001477***** |
| **Run Order 3** | **0.235922** | **0.04703** | **64.52479** | **5.016434635** | **4.39E-06***** |
| **Condition:Stimulation Time** | **-0.00124** | **0.00046** | **1875.815** | **-2.700214552** | **0.006992**** |

**Affective Stroop: Stimulation Time – Aversive Stimuli**

(Deconstruction: Stimulation Time)

| RT~Condition*Stimulation Time+Stimulation Site+Run Order+(1\|Subject) | | | | |  |
| --- | --- | --- | --- | --- | --- |
|  | Estimate | Error | df | t | Pr(>\|t\|) |
| **(Intercept)** | **0.593997** | **0.028687** | **29.16698** | **20.70610345** | **5.48E-19***** |
| **Condition** | **0.090189** | **0.009395** | **1848.945** | **9.599523566** | **2.50E-21***** |
| **Stimulation Time** | **-0.00747** | **0.001152** | **1871.616** | **-6.488695116** | **1.11E-10***** |
| Unguided dmPFC Site | 0.019174 | 0.030155 | 25.97822 | 0.635840322 | 0.530443 |
| **Run Order 2** | **0.123113** | **0.037623** | **34.15365** | **3.272306927** | **0.002444**** |
| **Run Order 3** | **0.217309** | **0.046514** | **68.97412** | **4.671921065** | **1.43E-05***** |
| Condition:Stimulation Time | -0.00041 | 0.000466 | 1849.052 | -0.881911551 | 0.377939 |

**Affective Stroop: Stimulation Time – Neutral Stimuli, Congruent Trials**

(Deconstruction: Stimulation Time)

| RT~Stimulation Time+Stimulation Site+Run Order+(1\|Subject) | | |  |  |  |
| --- | --- | --- | --- | --- | --- |
|  | Estimate | Error | df | t | Pr(>\|t\|) |
| **(Intercept)** | **0.581765** | **0.02798** | **28.57913** | **20.79204869** | **8.47E-19***** |
| **Stimulation Time** | **-0.00417** | **0.001448** | **948.9684** | **-2.881844958** | **0.004043**** |
| Unguided dmPFC Site | 0.019791 | 0.029504 | 25.82023 | 0.670781829 | 0.508316 |
| Run Order 2 | 0.063983 | 0.038427 | 39.94797 | 1.665047334 | 0.103728 |
| **Run Order 3** | **0.11364** | **0.051633** | **106.5662** | **2.200904003** | **0.0299*** |

**Affective Stroop: Stimulation time – Neutral Stimuli, Incongruent Trials**

(Deconstruction: Stimulation Time)

| RT~Stimulation Time+Stimulation Site+Run Order+(1\|Subject) | | | | | |
| --- | --- | --- | --- | --- | --- |
|  | Estimate | Error | df | t | Pr(>\|t\|) |
| **(Intercept)** | **0.664024** | **0.026145** | **29.6548** | **25.39732** | **1.12E-21***** |
| **Stimulation Time** | **-0.00699** | **0.001546** | **925.1762** | **-4.523317185** | **6.88E-06***** |
| Unguided dmPFC Site | 0.022207 | 0.02737 | 26.1225 | 0.811354926 | 0.424493 |
| **Run Order 2** | **0.113125** | **0.0369** | **45.38272** | **3.065687108** | **0.003651**** |
| **Run Order 3** | **0.213294** | **0.051786** | **134.0278** | **4.118793051** | **6.63E-05***** |

**Affective Stroop: Stimulation time – Negative Stimuli, Congruent Trials**

(Deconstruction: Stimulation Time)

| RT~Stimulation Time+Stimulation Site+Run Order+(1\|Subject) | | | | | |
| --- | --- | --- | --- | --- | --- |
|  | Estimate | Error | df | t | Pr(>\|t\|) |
| **(Intercept)** | **0.59813** | **0.030621** | **28.90028** | **19.53326198** | **3.39E-18***** |
| **Stimulation Time** | **-0.00726** | **0.00162** | **914.4986** | **-4.47979309** | **8.42E-06***** |
| Unguided dmPFC Site | 0.014715 | 0.032284 | 26.12064 | 0.455805399 | 0.652297 |
| **Run Order 2** | **0.113298** | **0.042353** | **41.33846** | **2.675052773** | **0.010657*** |
| **Run Order 3** | **0.210702** | **0.057204** | **110.7874** | **3.683375053** | **0.000358***** |

**Affective Stroop: Stimulation Time – Negative Stimuli, Incongruent Trials**

(Deconstruction: Stimulation Time)

| RT~Stimulation Time+Stimulation Site+Run Order+(1\|Subject) | | | | | |
| --- | --- | --- | --- | --- | --- |
|  | Estimate | Error | df | T | Pr(>\|t\|) |
| **(Intercept)** | **0.679348** | **0.0277** | **29.03046** | **24.52559195** | **5.91E-21***** |
| **Stimulation Time** | **-0.00772** | **0.001537** | **950.8891** | **-5.023017533** | **6.07E-07***** |
| Unguided dmPFC Site | 0.024098 | 0.029077 | 25.79715 | 0.828770825 | 0.414837 |
| **Run Order 2** | **0.127489** | **0.038615** | **42.73035** | **3.301549726** | **0.001948**** |
| **Run Order 3** | **0.214155** | **0.053139** | **121.2019** | **4.030087229** | **9.77E-05***** |

**Affective Stroop: Stimulation Time – Positive Stimuli, Congruent Trials**

(Deconstruction: Stimulation Time)

| RT~Stimulation Time+Stimulation Site+Run Order+(1\|Subject) | | | | | |
| --- | --- | --- | --- | --- | --- |
|  | Estimate | Error | df | t | Pr(>\|t\|) |
| **(Intercept)** | **0.585135** | **0.030212** | **28.72729** | **19.36743527** | **4.97E-18***** |
| **Stimulation Time** | **-0.00663** | **0.00154** | **923.5429** | **-4.306315675** | **1.84E-05***** |
| Unguided dmPFC Site | 0.006433 | 0.0319 | 26.08973 | 0.201650042 | 0.841754 |
| **Run Order 2** | **0.116382** | **0.0416** | **40.42387** | **2.797636628** | **0.007848**** |
| **Run Order 3** | **0.218999** | **0.055409** | **103.9702** | **3.952407084** | **0.000141***** |

**Affective Stroop: Stimulation Time – Positive Stimuli, Incongruent Trials**

(Deconstruction: Stimulation Time)

| RT~Stimulation Time+Stimulation Site+Run Order+(1\|Subject) | | | | | |
| --- | --- | --- | --- | --- | --- |
|  | Estimate | Error | df | t | Pr(>\|t\|) |
| **(Intercept)** | **0.675337** | **0.029731** | **29.16618** | **22.71488153** | **4.29E-20***** |
| **Stimulation Time** | **-0.00811** | **0.001568** | **972.6024** | **-5.172233957** | **2.81E-07***** |
| Unguided dmPFC Site | 0.012452 | 0.031292 | 26.17739 | 0.397941926 | 0.6939 |
| **Run Order 2** | **0.137261** | **0.040958** | **41.15585** | **3.351236772** | **0.001733**** |
| **Run Order 3** | **0.226918** | **0.055329** | **110.9504** | **4.101230192** | **7.86E-05***** |

**Affective Stroop: Accuracy**

(Omnibus: full factorial model testing Affective Stroop accuracy based on stimulation time)

| Correct~Condition*Valence*Stimulation Time+Stimulation Site+Run Order+(1\|Subject) | | | | |  |
| --- | --- | --- | --- | --- | --- |
|  | Estimate | Error | z | Pr(>\|z\|) | FDR |
| **(Intercept)** | **3.858659** | **0.530295** | **7.276433** | **3.43E-13***** | **6.52E-12** |
| **Condition** | **-0.8829** | **0.524914** | **-1.68199** | **0.092571346^.^** | **0.293143** |
| Aversive | -0.56303 | 0.549019 | -1.02552 | 0.305118442 | 0.527023 |
| Pleasant | -0.77154 | 0.566967 | -1.36081 | 0.173572311 | 0.394361 |
| **Stimulation Time** | **0.087167** | **0.032522** | **2.680219** | **0.00735741**** | **0.034948** |
| Unguided dmPFC Site | 0.403286 | 0.320886 | 1.256787 | 0.208830663 | 0.396778 |
| **Run Order 2** | **-1.5095** | **0.48769** | **-3.0952** | **0.001966812**** | **0.012456** |
| **Run Order 3** | **-2.94425** | **0.777402** | **-3.7873** | **0.000152296***** | **0.001447** |
| Condition:Aversive | 0.344216 | 0.673362 | 0.511191 | 0.609217649 | 0.759328 |
| Condition:Pleasant | 0.446004 | 0.683749 | 0.652291 | 0.514213198 | 0.751542 |
| Condition:Stimulation Time | -0.00695 | 0.024397 | -0.28501 | 0.775639784 | 0.818731 |
| Aversive:Stimulation Time | -0.01059 | 0.025435 | -0.4163 | 0.677193832 | 0.759328 |
| Pleasant:Stimulation Time | 0.037214 | 0.02819 | 1.320097 | 0.186802756 | 0.394361 |
| Condition:Aversive:Stimulation Time | 0.012864 | 0.031126 | 0.413284 | 0.679398541 | 0.759328 |
| Condition:Pleasant:Stimulation Time | -0.03184 | 0.033337 | -0.95502 | 0.339569492 | 0.537652 |

**Affective Stroop: Accuracy**

(Omnibus; Relevel: Aversive stimuli)

| Correct~Condition*Valence*Stimulation Time+Stimulation Site+Run Order+(1\|Subject) | | | | |  |
| --- | --- | --- | --- | --- | --- |
|  | Estimate | Error | z | Pr(>\|z\|) | FDR |
| **(Intercept)** | **3.295348** | **0.442699** | **7.44376** | **9.79E-14***** | N/A |
| Condition | -0.53918 | 0.421415 | -1.27944 | 0.200741097 | N/A |
| Neutral | 0.564777 | 0.549232 | 1.028303 | 0.303807299 | N/A |
| Pleasant | -0.20919 | 0.48455 | -0.43171 | 0.665950554 | 0.759328 |
| **Stimulation Time** | **0.076678** | **0.029506** | **2.598754** | **0.009356268**** | N/A |
| Unguided dmPFC Site | 0.403738 | 0.320987 | 1.257802 | 0.208463342 | N/A |
| **Run Order 2** | **-1.51063** | **0.487729** | **-3.09728** | **0.001953085*** | N/A |
| **Run Order 3** | **-2.94677** | **0.777428** | **-3.7904** | **0.000150403***** | N/A |
| Condition:Neutral | -0.34585 | 0.673556 | -0.51347 | 0.607620778 | N/A |
| Condition:Pleasant | 0.102915 | 0.607979 | 0.169275 | 0.865580588 | 0.865581 |
| Condition:Stimulation Time | 0.005924 | 0.019304 | 0.306854 | 0.758954566 | N/A |
| Neutral:Stimulation Time | 0.01051 | 0.025441 | 0.413125 | 0.679515368 | N/A |
| **Pleasant:Stimulation Time** | **0.047826** | **0.024492** | **1.952678** | **0.050857744^.^** | **0.193259** |
| Condition:Neutral:Stimulation Time | -0.01278 | 0.031131 | -0.4105 | 0.681439903 | N/A |
| Condition:Pleasant:Stimulation Time | -0.04474 | 0.029837 | -1.49943 | 0.133762873 | 0.363071 |

**Affective Stroop: Accuracy – Neutral Trials**

(Deconstruction: Accuracy)

| Correct~Condition*Stimulation Time+Stimulation Site+Run Order+(1\|Subject) | | | | |
| --- | --- | --- | --- | --- |
|  | Estimate | Error | z | Pr(>\|z\|) |
| **(Intercept)** | **3.973973** | **0.567814** | **6.998723** | **2.58E-12***** |
| Condition | -0.82719 | 0.523019 | -1.58158 | 0.113746 |
| Stimulation Time | 0.07165 | 0.048827 | 1.467419 | 0.142262 |
| Unguided dmPFC Site | 0.294894 | 0.393023 | 0.750322 | 0.453061 |
| **Run Order 2** | **-1.43666** | **0.722055** | **-1.98969** | **0.046625*** |
| **Run Order 3** | **-2.59434** | **1.288628** | **-2.01326** | **0.044087*** |
| Condition:Stimulation Time | -0.00747 | 0.024268 | -0.30796 | 0.758115 |

**Affective Stroop: Accuracy – Aversive Trials**

(Deconstruction: Accuracy)

| Correct~Condition*Stimulation Time+Stimulation Site+Run Order+(1\|Subject) | | | | |
| --- | --- | --- | --- | --- |
|  | Estimate | Error | z | Pr(>\|z\|) |
| **(Intercept)** | **3.158682** | **0.461648** | **6.842192** | **7.80E-12***** |
| Condition | -0.57308 | 0.414976 | -1.38099 | 0.167283347 |
| **Stimulation Time** | **0.104449** | **0.04207** | **2.48275** | **0.013037245**** |
| **Unguided dmPFC Site** | **0.786677** | **0.365546** | **2.152059** | **0.031392728*** |
| **Run Order 2** | **-1.91807** | **0.646923** | **-2.96491** | **0.003027671**** |
| **Run Order 3** | **-3.85956** | **1.14625** | **-3.36712** | **0.000759575***** |
| Condition:Stimulation Time | 0.007783 | 0.018962 | 0.410435 | 0.681486753 |

**Affective Stroop: Accuracy – Pleasant Trials**

(Deconstruction: Accuracy)

| Correct~Condition*Stimulation Time+Stimulation Site+Run Order+(1\|Subject) | | | | |
| --- | --- | --- | --- | --- |
|  | Estimate | Error | z | Pr(>\|z\|) |
| **(Intercept)** | **3.160811** | **0.470568** | **6.717017** | **1.85E-11***** |
| Condition | -0.49144 | 0.437269 | -1.12389 | 0.261059727 |
| **Stimulation Time** | **0.095418** | **0.045022** | **2.119348** | **0.034061033*** |
| Unguided dmPFC Site | 0.263752 | 0.356755 | 0.739308 | 0.459720093 |
| **Run Order 2** | **-1.12549** | **0.650556** | **-1.73004** | **0.083623176^.^** |
| **Run Order 3** | **-2.09625** | **1.162721** | **-1.80288** | **0.071407007^.^** |
| Condition:Stimulation Time | -0.03618 | 0.022754 | -1.59006 | 0.111821783 |

**Affective Stroop: Accuracy – Condition, Neutral Trials**

(Deconstruction: Accuracy)

| Correct~Stimulation Time+Stimulation Site+Run Order+(1\|Subject) | | | |  |
| --- | --- | --- | --- | --- |
|  | Estimate | Error | z | Pr(>\|z\|) |
| **(Intercept)** | **3.900894** | **0.809625** | **4.818148** | **1.45E-06***** |
| **Stimulation Time** | **0.142922** | **0.086737** | **1.647767** | **0.099400456^.^** |
| Unguided dmPFC Site | 0.564563 | 0.682992 | 0.826603 | 0.408462225 |
| Run Order 2 | -2.15646 | 1.320275 | -1.63334 | 0.102396615 |
| **Run Order 3** | **-4.71206** | **2.449178** | **-1.92394** | **0.054362453*** |

**Affective Stroop: Accuracy – Incongruent, Neutral Trials**

(Deconstruction: Accuracy)

| Correct~Stimulation Time+Stimulation Site+Run Order+(1\|Subject) | | | |  |
| --- | --- | --- | --- | --- |
|  | Estimate | Error | z | Pr(>\|z\|) |
| **(Intercept)** | **3.187098** | **0.437067** | **7.292016** | **3.05E-13***** |
| Stimulation Time | 0.034662 | 0.051077 | 0.678624 | 0.497376338 |
| Unguided dmPFC Site | 0.127439 | 0.365728 | 0.348453 | 0.727500142 |
| Run Order 2 | -1.09625 | 0.756354 | -1.44939 | 0.1472299 |
| Run Order 3 | -1.7346 | 1.404923 | -1.23466 | 0.216956672 |

**Affective Stroop: Accuracy – Congruent, Positive Trials**

(Deconstruction: Accuracy)

| Correct~Stimulation Time+Stimulation Site+Run Order+(1\|Subject) | | | |  |
| --- | --- | --- | --- | --- |
|  | Estimate | Error | z | Pr(>\|z\|) |
| **(Intercept)** | **3.591744** | **0.766113** | **4.68827** | **2.76E-06***** |
| Stimulation Time | 0.124726 | 0.087618 | 1.423518 | 0.154586149 |
| Unguided dmPFC Site | 0.33977 | 0.73971 | 0.459329 | 0.645998044 |
| Run Order 2 | -1.86417 | 1.321459 | -1.41069 | 0.158335484 |
| Run Order 3 | -2.65504 | 2.410132 | -1.10162 | 0.270628893 |

**Affective Stroop: Accuracy – Incongruent, Positive Trials**

(Deconstruction: Accuracy)

| Correct~Stimulation Time+Stimulation Site+Run Order+(1\|Subject) | | | |  |
| --- | --- | --- | --- | --- |
|  | Estimate | Error | z | Pr(>\|z\|) |
| **(Intercept)** | **2.518976** | **0.360752** | **6.982574** | **2.90E-12***** |
| Stimulation Time | 0.050867 | 0.045146 | 1.126738 | 0.259853328 |
| Unguided dmPFC Site | 0.293388 | 0.31458 | 0.932635 | 0.351008439 |
| Run Order 2 | -0.91023 | 0.656037 | -1.38746 | 0.165300985 |
| Run Order 3 | -1.87039 | 1.231681 | -1.51857 | 0.128870954 |

**Affective Stroop: Accuracy – Congruent, Unpleasant Trials**

(Deconstruction: Accuracy)

| Correct~Stimulation Time+Stimulation Site+Run Order+(1\|Subject) | | |  |  |
| --- | --- | --- | --- | --- |
|  | Estimate | Std.Error | z | Pr(>\|z\|) |
| **(Intercept)** | **3.187537** | **0.542802** | **5.872379** | **4.30E-09***** |
| **Stimulation Time** | **0.202212** | **0.061428** | **3.291886** | **0.000995***** |
| **Unguided dmPFC Site** | **0.900513** | **0.432578** | **2.081737** | **0.037367*** |
| **Run Order 2** | **-3.71819** | **0.92299** | **-4.02842** | **5.62E-05***** |
| **Run Order 3** | **-6.85122** | **1.733438** | **-3.95239** | **7.74E-05***** |

**Affective Stroop: Accuracy – Incongruent, Unpleasant Trials**

(Deconstruction: Accuracy)

| Correct~Stimulation Time+Stimulation Site+Run Order+(1\|Subject) | | | |  |
| --- | --- | --- | --- | --- |
|  | Estimate | Error | z | Pr(>\|z\|) |
| **(Intercept)** | **2.668485** | **0.462105** | **5.774624** | **7.71E-09***** |
| Stimulation Time | 0.048947 | 0.051647 | 0.947724 | 0.34327008 |
| **Unguided dmPFC Site** | **0.778393** | **0.438796** | **1.77393** | **0.076074696^.^** |
| Run Order 2 | -0.81134 | 0.811104 | -1.00029 | 0.317171769 |
| Run Order 3 | -2.04133 | 1.454619 | -1.40335 | 0.160513629 |

**Color-Word Stroop: Response Time (Trial)**

(Omnibus: Model testing Color-Word Stroop response time based on trial)

| RT~Trials*Condition+(1\|Subject) | Estimate | Std.Error | df | t | Pr(>\|t\|) |
| --- | --- | --- | --- | --- | --- |
| **(Intercept)** | **0.717121** | **0.023637** | **31.26825** | **30.33917** | **9.24E-25***** |
| Trials | 2.53E-05 | 2.48E-05 | 9445.352 | 1.022364 | 0.306635 |
| **Condition** | **0.073361** | **0.006922** | **9440.028** | **10.59793** | **4.27E-26***** |
| Trials:Condition | -2.58E-05 | 3.45E-05 | 9440.03 | -0.74878 | 0.45401 |

**Color-Word Stroop: Correct (Trial)**

(Omnibus: Model testing Color-Word Stroop response time based on trial)

| Correct~Condition*Trials+(1\|Sub) | Estimate | Std.Error | z | Pr(>\|z\|) |
| --- | --- | --- | --- | --- |
| **(Intercept)** | **3.809365** | **0.216315** | **17.61026** | **2.06E-69***** |
| **Condition** | **-0.50813** | **0.208753** | **-2.43414** | **0.014927**** |
| Trials | -0.00075 | 0.000762 | -0.97899 | 0.327583 |
| Condition:Trials | 0.001376 | 0.001017 | 1.352243 | 0.176298 |

**Color-Word Stroop: Stimulation Time (Trial)**

(Omnibus: Full-factorial model testing Color-Word Stroop response time based on trial and stimulation time)

| RT~Condition*Stimulation Time*Stimulation Site+Run Order +Trials+(1\|Sub) | | | | | |
| --- | --- | --- | --- | --- | --- |
|  | Estimate | Std.Error | df | t | Pr(>\|t\|) |
| **(Intercept)** | **0.643608** | **0.046414** | **28.52468** | **13.86665** | **3.33E-14***** |
| **Condition** | **0.060014** | **0.010248** | **9431.869** | **5.856112** | **4.90E-09***** |
| Stimulation Time | 0.001856 | 0.001312 | 7756.88 | 1.414789 | 0.15717 |
| **rmPFC Site** | **0.031128** | **0.011064** | **9439.061** | **2.813501** | **0.004911**** |
| **Unguided dmPFC Site** | **0.171392** | **0.05796** | **36.37361** | **2.957056** | **0.005428**** |
| Run Order 2 | 0.012658 | 0.060932 | 27.90664 | 0.207743 | 0.836939 |
| Run Order 3 | 0.083075 | 0.066092 | 34.38033 | 1.256949 | 0.217245 |
| **Trials** | **5.95E-05** | **2.30E-05** | **9456.261** | **2.584737** | **0.00976**** |
| **Condition:Stimulation Time** | **0.001362** | **0.000588** | **9431.876** | **2.316996** | **0.020525*** |
| Condition:rmPFC Site | 0.008917 | 0.015316 | 9431.897 | 0.582195 | 0.560449 |
| Condition:Unguided dmPFC Site | 0.01538 | 0.016097 | 9431.866 | 0.955473 | 0.339363 |
| Stimulation Time:rmPFC Site | -0.00117 | 0.000611 | 9436.782 | -1.92042 | 0.054835 |
| **Stimulation Time:Unguided dmPFC Site** | **-0.01016** | **0.001667** | **3476.269** | **-6.09427** | **1.22E-09***** |
| Condition:Stimulation Time:rmPFC Site | -0.00125 | 0.000853 | 9431.908 | -1.46166 | 0.143869 |
| **Condition:Stimulation Time:Unguided dmPFC Site** | **-0.00293** | **0.000926** | **9431.865** | **-3.1603** | **0.001581**** |

**Color-Word Stroop: Accuracy (Trial)**

(Omnibus: Full-factorial model testing Color-Word Stroop response time based on trial and stimulation time)

| Correct~Condition*Stimulation Time*Stimulation Site+Run Order +Trials+(1\|Sub) | | | | |
| --- | --- | --- | --- | --- |
|  | Estimate | Std.Error | z | Pr(>\|z\|) |
| **(Intercept)** | **4.005922** | **0.387731** | **10.33171** | **5.06E-25***** |
| **Condition** | **-0.71306** | **0.309472** | **-2.30413** | **0.021215*** |
| **Stimulation Time** | **-0.10361** | **0.032941** | **-3.14537** | **0.001659**** |
| rmPFC Site | 0.35181 | 0.375682 | 0.936455 | 0.349039 |
| Unguided dmPFC Site | -0.61217 | 0.580363 | -1.0548 | 0.291517 |
| **Run Order 2** | **1.09378** | **0.490067** | **2.231896** | **0.025622*** |
| **Run Order 3** | **1.786211** | **0.76127** | **2.346356** | **0.018958*** |
| Trials | 0.000866 | 0.000687 | 1.260153 | 0.207614 |
| **Condition:Stimulation Time** | **0.03706** | **0.017534** | **2.113567** | **0.034552*** |
| Condition:rmPFC Site | -0.46816 | 0.458646 | -1.02075 | 0.307374 |
| Condition:Unguided dmPFC Site | 0.196023 | 0.460021 | 0.426117 | 0.670022 |
| Stimulation Time:rmPFC Site | -0.01961 | 0.019249 | -1.0186 | 0.308392 |
| **Stimulation Time:Unguided dmPFC Site** | **0.072716** | **0.033678** | **2.15919** | **0.030835*** |
| Condition:Stimulation Time:rmPFC Site | 0.023519 | 0.025233 | 0.932081 | 0.351295 |
| Condition:Stimulation Time:Unguided dmPFC Site | -0.02912 | 0.029942 | -0.97267 | 0.330716 |

**Affective Stroop: Correct (Trial)**

(Omnibus: Model testing Affective Stroop Response Time based on trial)

| Correct~Condition*Trials+(1\|Sub) | Estimate | Std.Error | z | Pr(>\|z\|) |
| --- | --- | --- | --- | --- |
| **(Intercept)** | **3.045222** | **0.23671** | **12.86476** | **7.11E-38***** |
| **Condition** | **-0.4169** | **0.229752** | **-1.81459** | **0.069587^.^** |
| **Trials** | **0.006184** | **0.001856** | **3.332166** | **0.000862***** |
| **Condition:Trials** | **-0.00444** | **0.00224** | **-1.98164** | **0.047519*** |

**Affective Stroop: Response Time (Trial)**

(Omnibus: Model testing Affective Stroop Response Time based on trial)

| RT~Trials*Condition+(1\|Sub) | Estimate | Std.Error | df | t | Pr(>\|t\|) |
| --- | --- | --- | --- | --- | --- |
| **(Intercept)** | **0.612089** | **0.013304** | **34.25316** | **46.00777** | **2.20E-32***** |
| **Trials** | **-0.00026** | **3.50E-05** | **5655.057** | **-7.43848** | **1.17E-13***** |
| **Condition** | **0.087375** | **0.005622** | **5655.047** | **15.5421** | **2.26E-53***** |
| Trials:Condition | -5.54E-05 | 4.90E-05 | 5655.069 | -1.13016 | 0.258457 |

**Affective Stroop: Stimulation Time (Trial)**

(Omnibus: Full-factorial model testing Affective Stroop response time based on stimulation time and including trial)

|  | Estimate | Std.Error | df | t | Pr(>\|t\|) |
| --- | --- | --- | --- | --- | --- |
| **(Intercept)** | **0.594565** | **0.022823** | **35.50135** | **26.05156** | **9.89E-25***** |
| **Condition** | **0.075162** | **0.00917** | **5647.138** | **8.196515** | **3.03E-16***** |
| Stimulation Time | 0.000314 | 0.001191 | 38.68741 | 0.263749 | 0.793373 |
| Aversive | 0.005031 | 0.009306 | 5647.411 | 0.540595 | 0.588808 |
| Pleasant | -0.01152 | 0.009229 | 5647.242 | -1.24822 | 0.212003 |
| **Trials** | **-0.00031** | **4.99E-05** | **59.19718** | **-6.2986** | **4.06E-08***** |
| Unguided dmPFC | 0.027939 | 0.026521 | 27.02728 | 1.053469 | 0.301454 |
| Condition:Stimulation Time | 0.000598 | 0.000455 | 5647.324 | 1.313922 | 0.188926 |
| Condition:Aversive | 0.015476 | 0.013008 | 5647.38 | 1.189769 | 0.234187 |
| Condition:Pleasant | **0.02431** | **0.012996** | **5647.291** | **1.870566** | **0.061457^.^** |
| Stimulation Time:Aversive | 0.000389 | 0.00046 | 5647.233 | 0.844194 | 0.398597 |
| **Stimulation Time:Pleasant** | **0.00128** | **0.000454** | **5647.156** | **2.819624** | **0.004825**** |
| **Condition:Stimulation Time:Aversive** | **-0.00108** | **0.000646** | **5647.448** | **-1.66766** | **0.095438^.^** |
| **Condition:Stimulation Time:Pleasant** | **-0.00188** | **0.000644** | **5647.513** | **-2.92099** | **0.003503**** |

**Affective Stroop: Accuracy (Trial)**

(Omnibus: Full-factorial model testing Affective Stroop accuracy based on stimulation time and including trial)

| Correct~Condition*Valence*Stimulation Time+Stimulation Site+Trials+Run Order +(1\|Sub) | | | | |
| --- | --- | --- | --- | --- |
|  | Estimate | Std.Error | z | Pr(>\|z\|) |
| **(Intercept)** | **3.84922** | **0.53128** | **7.245179** | **4.32E-13***** |
| Condition | -0.88222 | 0.524695 | -1.68139 | 0.092687 |
| Aversive | -0.56298 | 0.548883 | -1.02568 | 0.305042 |
| Pleasant | -0.77353 | 0.566758 | -1.36484 | 0.172304 |
| Stimulation Time | 0.108173 | 0.078955 | 1.370055 | 0.17067 |
| Unguided dmPFC | 0.444914 | 0.351572 | 1.265497 | 0.205693 |
| Trials | -0.00089 | 0.003058 | -0.29226 | 0.77009 |
| **Run Order 2** | **-1.76439** | **1.000389** | **-1.7637** | **0.077782^.^** |
| **Run Order 3** | **-3.49228** | **2.032083** | **-1.71857** | **0.085693^.^** |
| Condition:Aversive | 0.341642 | 0.673187 | 0.507499 | 0.611805 |
| Condition:Pleasant | 0.447588 | 0.683519 | 0.654829 | 0.512578 |
| Condition:Stimulation Time | -0.00699 | 0.024389 | -0.28663 | 0.774398 |
| Aversive:Stimulation Time | -0.01058 | 0.02543 | -0.41617 | 0.677287 |
| Pleasant:Stimulation Time | 0.037299 | 0.028186 | 1.323315 | 0.185731 |
| Condition:Aversive:Stimulation Time | 0.012963 | 0.031119 | 0.416578 | 0.676987 |
| Condition:Pleasant:Stimulation Time | -0.03191 | 0.033332 | -0.95735 | 0.338392 |
